# Multi-omics analysis of keratinocytes reveals dermokine-dependent regulation of cell-cell adhesion via p120

**DOI:** 10.1101/2025.01.10.632346

**Authors:** Vahap Canbay, Till Wüstemann, Weihua Tian, Tobias A. Beyer, Camilla Reiter Elbæk, Michael Stumpe, Gaetana Restivo, Chatpakorn Christiansen, Anabel Migenda Herranz, Susanne Mailand, Jürg Hafner, Rune Busk Damgaard, Steffen Goletz, Jörn Dengjel, Ulrich auf dem Keller, Chiara Francavilla

## Abstract

Loss of keratinocyte differentiation is a leading cause in several skin diseases and needs to be controlled in adult homeostasis by for instance growth factors and proteases. Among them, we studied the role of isoform-rich dermokine – a wound- and tumour-related matrix metalloproteinase 10 substrate – via functional multi-omics. We generated dermokine isoform-dependent keratinocyte knockouts and three dimensional (3D) organotypic skin cultures and analyzed changes in their proteome and phosphoproteome by quantitative mass spectrometry. Through functional *in vitro* assays, we demonstrate that in the absence of dermokine-isoforms, p120 phosphorylation increases while cell-cell adhesion decreases in keratinocytes. Furthermore, we validate the link between decreased dermokine expression and phosphorylated p120-mediated adhesion in non-healing wounds samples derived from patients. Our data reveal a novel dermokine-p120-dependent cell-cell adhesion phenotype in keratinocytes and improve our understanding of wound-edge keratinocytes, expanding the hypothesis that dysregulated wounds resemble cancer.

## Introduction

The skin is the largest organ of the human body and is composed of epidermis, dermis and subcutaneous layer^1^. Keratinocytes, the most prominent cell type of the epidermis, is forming a stratified epithelial through terminal differentiation from cells of the basal to the spinous, granular and corneal layers^2^. Through this stratified epithelium, the epidermis serves as a protective barrier against external threats, water loss, and mechanical strains^1^. Loss or aberrant keratinocyte differentiation is causative to a plethora of skin diseases, such as psoriasis, cutaneous squamous cell carcinoma (SCC) and non-healing wounds^3^. In SCC keratinocytes lose the ability to differentiate into a stratified epithelium and undergo epithelial to mesenchymal transformation (EMT) mediated by the loss of cell-cell adhesion proteins^4^. In psoriatic tissue, keratinocytes are exposed to sustained inflammation causing excessive proliferation, and aberrant differentiation^5^. Further, in the context of non-healing wounds, keratinocytes at the rim of non-healing wounds remain in a vicious cycle of chronic inflammation. This prevents keratinocyte to re-epithelialize the wound bed by proliferation and migration^6–8^. In summary, keratinocyte differentiation must be tightly regulated in adult skin homeostasis to prevent such devastating diseases. Therefore, a deeper understanding of the molecular determinants underlying keratinocytes differentiation would open novel avenues for the treatment of skin diseases.

Several players have been described regulating keratinocyte differentiation, including growth factors, calcium homeostasis, and proteases^9,10^. Specifically, changes in the abundance, activity, substrate availability as well as inhibitory mechanisms of proteases have been associated to keratinocytes differentiation. Among them are kallikrein-related peptides (KLK), the desquamation cascade and the cathepsin related activation of transglutaminases (TGM)^11^. One of the major functions of proteases is the sequential cleavage of proteins like pro-filaggrin (pro-FLG), epidermal growth factor (EGF), and G-protein coupled receptors thereby regulating their activities in the skin. Overall, studying the proteases substrate repertoire is key to understand keratinocytes differentiation^9^. Among proteases with a function in keratinocytes, the wound and cancer-associated matrix metalloproteinase 10 (MMP10), which is expressed in wound-edge keratinocytes of skin lesions, plays a crucial role in the cleavage of proteins regulating keratinocytes migration and adhesion. We have previously identified dermokine, a member of the stratified epithelium-secreted peptides complex, as a novel substrate of MMP10 in basal keratinocytes^12^. Dermokine is gradually expressed by differentiating keratinocytes, absent in the basal layer, minimally present in the spinous layer, and most abundant in the granular layer^13^. Even though a putative role of dermokine in modulating keratinocyte differentiation has been suggested in mice^14^, the precise molecular mechanism underlying dermokine functions in human keratinocytes has not yet been described yet.

Dermokine is encoded by the *DMKN* gene which produces multiple transcripts resulting in three major protein isoforms: dermokine-α, -β, and -γ^15^. Dermokine-β and -γ are secreted into the extracellular space and found in the granular layers of the epidermis whereas dermokine-α is expressed equally among all epidermal layers^13^. In murine models, the depletion of dermokine-βγ results in severe conditions like neonatal skin hyperkeratosis, resembling psoriasis-like skin disorders^14^. As mice deficient in all dermokine isoforms show a more severe phenotype, a compensatory role for dermokine-α has been suggested during keratinocytes differentiation^14^. Here, to reveal the role of dermokine in keratinocyte differentiation in the human context we used CRISPR/Cas9-mediated depletion of dermokine isoforms in human keratinocytes using three-dimensional (3D) organotypic skin cultures as model^16,17^. These 3D cultures resemble epidermis-like structures, including differentiating keratinocytes and have been used as an experimental model to study, for example, the invasiveness of keratinocyte-derived SCC^18,19^. To characterize the functions of DMKN in human epidermal differentiation, these genetically engineered 3D cultures were subjected to combined omics approaches followed by functional assays. By targeted and quantitative proteomics, we showed that dermokine affected epidermal proteins expression pattern in 3D cultures high lightening the role of dermokine in keratinocyte differentiation. Subsequently, we used cutting-edge data-independent acquisition-based (DIA) phosphoproteomics, functional *in vitro* assays, and staining of patient biopsies from healthy and pathological skin to reveal changes in cell-cell adhesion proteins downstream of dermokine. Specifically, the phosphorylation pattern of p120, a known regulator of cell-cell adhesion^20^ was increased upon ablation of dermokine isoforms. In summary, multi-omics, functional analyses of keratinocyte models and skin tissue staining reveal that dermokine regulates cell-cell adhesion of keratinocytes through p120 *in vitro* and that dermokine expression as well as p120 phosphorylation were dysregulated in non-healing wounds *in vivo*.

## Results

### Dermokine regulates the thickness of epidermis-like structure and the proteome of keratinocytes

To study the role of dermokine in keratinocyte differentiation^12^we edited the genomic sites encoding for either dermokine-βγ or -αβ (*DMKN* βγ or αβ) in human keratinocytes by CRISPR/Cas9 technology^16,21^ (Figure 1A). We used two CRISPR/Cas9-based methods to generate independent dermokine knockouts (KO) in keratinocytes (Figures 1B-1C). The first method, “chemical transfection”, was based on fluorescence-activated cell sorting to select single-cell clones, resulting in *DMKN* βγ^−/−^ clone^22^ (Figure 1B). The second method, “FluidFM™ multiplexed CRISPR approach” was based on co-injecting CRISPR ribonucleoprotein (RNP) complexes and green fluorescent protein (GFP) mRNA into keratinocytes, thus eliminating exon 17 and 1 of dermokine-β and -α respectively, resulting in *DMKN* αβ^−/−^ keratinocytes^21^ (Figure 1C). We confirmed the KO of *DMKN* genes generated with either method by Sanger sequencing (Figure S1A-D). We then grew 3D organotypic skin cultures using keratinocytes either from WT or from the *DMKN* βγ^−/−^ or *DMKN* αβ^−/−^ clones on top of primary human fibroblasts incorporated within matrigel^16^ (Figures 1B-C). To confirm changes in the expression of dermokine isoforms in this model system we used targeted proteomics and synthetic reference peptides^23^. We selected five proteotypic peptides spread across the entire dermokine sequence, including two near the N-terminus, two after the keratin-like domain, and one at the C-terminus truncated (Figure 1D). We calculated the summed transition areas of all proteotypic peptides in either *DMKN* βγ^−/−^ or *DMKN* αβ^−/−^ relative to WT keratinocytes and confirmed a substantial decreased in dermokine protein abundance (Figures S1E-S1F). We focused on the C-terminal peptide - GGVSPSSSASR -, since it was shared by both dermokine-β and -α (Fig 1D). If only dermokine-β was truncated, leaving the -α isoform intact, this peptide would still be detectable. We observed only a slight reduction in the peptide peak area when dermokine-β, in combination with -γ, was truncated (Figure 1E). However, upon removing both dermokine-α and -β, sharing the cytokine-like domain where the C-terminal peptide was located, we observed a substantial twofold decrease in the abundance of the truncated peptide (Figure 1F). Overall, this data confirms the absence of *DMKN* at the protein level.

**Figure 1:**
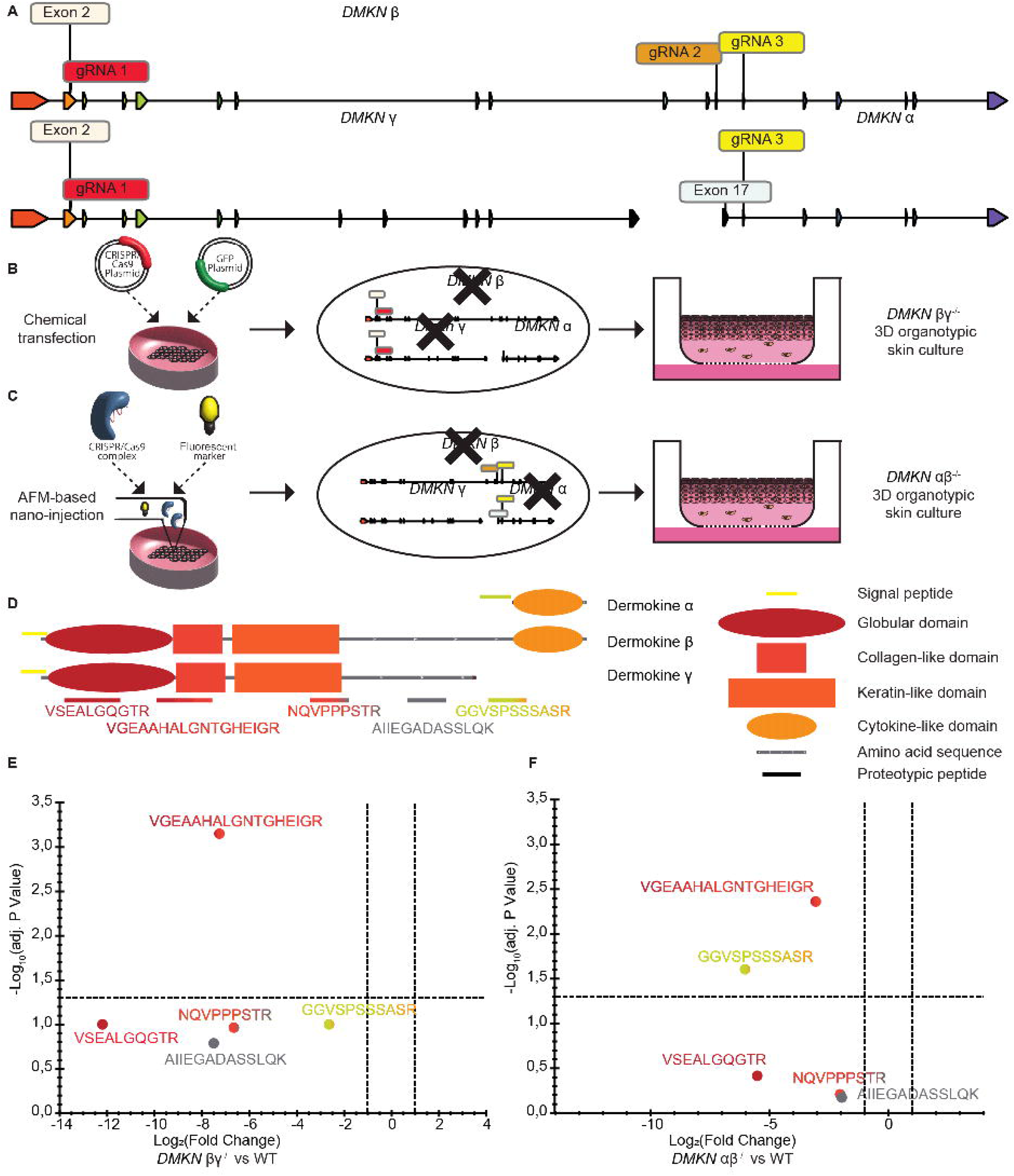
Targeted proteomics validates dermokine-ablated human keratinocytes. **(A)** Schematic of the genetic loci of the dermokine isoforms and locations of guide RNAs (gRNA 1, gRNA 2, gRNA 3) targeting isoform specific exons. **(B and C)** Methodologies used to delete dermokine in keratinocytes through either “chemical transfection” **(B)** or “FluidFM™ multiplexed CRISPR approach”. **(C)** After transfection and generation of monoclonal cell lines, WT, *DMKN* αβ^−/−^ or *DMKN* βγ^−/−^ keratinocytes were placed in 3D organotypic skin cultures. **(D)** Schematic of the protein domains of the dermokine isoforms and of the five isoform-specific and proteotypic peptides used for targeted proteomics analysis. **(E and F)** Proteotypic dermokine peptide abundances are Log_2_ transformed fold changes of summed transition areas from *DMKN* βγ^−/−^ (**E**) or *DMKN* αβ^−/−^ **(F)** relative to WT keratinocytes analysed by targeted proteomics. N = 3 different 3D organotypic skin cultures.

To study how the *DMKN* KO genotype influenced the phenotype of our model system we analysed WT and *DMKN* KO 3D cultures by immunohistochemical (IHC) staining and confirmed the lack of dermokine expression in both *DMKN* βγ^−/−^ and *DMKN* αβ^−/−^ (Figure 2). IHC staining of the *DMKN* KO samples revealed a thinner total epidermis-like structure, with keratinocytes packed tightly together and the lack of extension of the suprabasal layers of keratinocytes beyond the spinous layer. This observation was consistent with the known dermokine expression pattern in keratinocytes^13^. Next, we evaluated whether the lack of dermokine affected the early-stage differentiation marker by staining for keratin 14 (KRT14) and suprabasal keratin 10 (KRT10)^1^. Both *DMKN* KOs showed an expression of KRT14 comparable to the WT, whereas *DMKN* βγ^−/−^ but not *DMKN* αβ^−/−^, decreased the expression of KRT10 compared to WT (Figure 2). The presence of filaggrin (FLG), which is a marker of the granular layer in keratinocytes^1^, was confirmed in the WT keratinocytes but not observed in the *DMKN* KO keratinocytes (Figure 2). This finding suggests that the dermokine isoforms indirectly decreases the expression of late-stage keratinocyte proteins. The expression of ITGα6, which is a marker of the basal layer, was similar in WT and *DMKN* KO keratinocytes, confirming that the expression of dermokine does not affect the adhesion of basal keratinocytes to the underlying matrix^1^ (Figure 2). EGF receptor (EGFR) expressed in the early differentiating layer, indicative of proliferative keratinocytes, showed only a slight decreased expression in *DMKN* KOs compared to WT cells^24^ (Figure 2). In conclusion, our data shows that dermokine regulates epidermal thickness, confirming the phenotype of murine models^14^.

**Figure 2:**
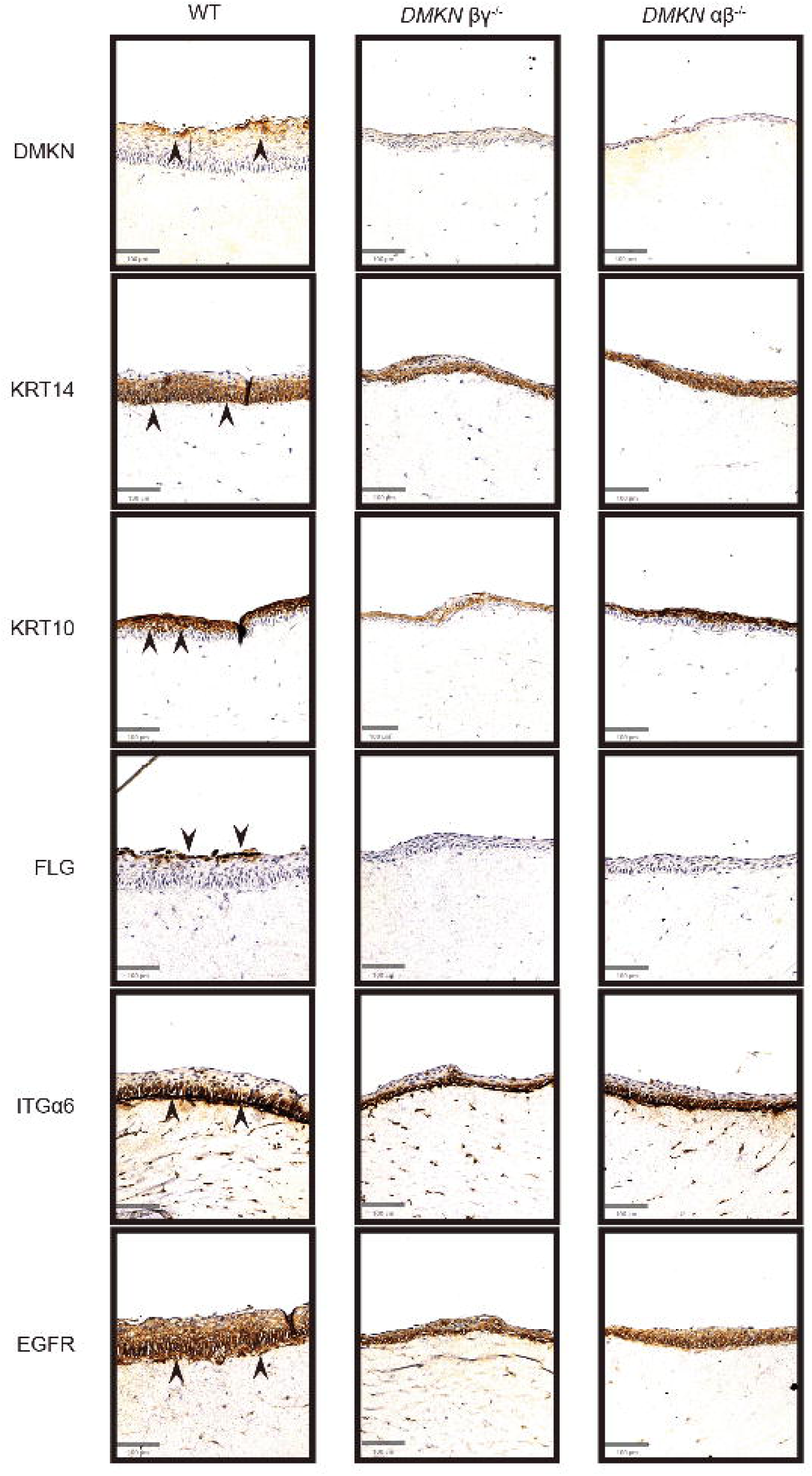
Immunohistochemical staining of WT, DMKN αβ^−/−^ or DMKN βγ^−/−^ 3D organotypic skin cultures tissue shows changes in epithelial markers. Immunohistochemical staining with the indicated antibodies of WT and *DMKN* αβ^−/−^ or *DMKN* βγ^−/−^ keratinocytes 3D organotypic skin cultures. N = 3, scale bar = 100 µm.

To further characterize the phenotype of *DMKN* KOs in an unbiased manner, we analyzed changes in the proteome of the 3D cultures grown from either WT or *DMKN* KO keratinocytes via LC-MS-based proteomics. In total, we quantified 3753 proteins in WT, *DMKN* βγ^−/−^ and *DMKN* αβ^−/−^ 3D cultures with a coefficient of variation (CV) of less than 15 % between the replicates (Figure S2A). The Pearson correlation analysis of WT, *DMKN* βγ^−/−^ and *DMKN* αβ^−/−^ 3D cultures revealed high correlation among replicates for each experimental condition (Figure S2B), in line with previous publications^25,26^. Finally, principal component analysis (PCA) demonstrated that the proteome of the three conditions was well separated (Figure S3C). Hierarchical clustering revealed clusters including proteins more abundant only in the WT (Cluster 2), in both the KO (Cluster 4), and only in one of the two KO samples (Clusters 1 and 3) (Figure S2D), which suggests that the two dermokine isoforms may have overlapping functions in keratinocytes. GO term enrichment analysis showed that cluster 2 is linked to terms like “formation of the cornified envelope” and cluster 4 associated with signalling pathways (Figure S2E). The differential abundance analysis, visualized by volcano plot, revealed 312 and 231 significantly increased proteins in the *DMKN* βγ^−/−^ and *DMKN* αβ^−/−^ 3D cultures, respectively, when compared to the WT (Figure 3A). We also identified 217 and 230 significantly decreased proteins in the respective *DMKN* βγ^−/−^ and *DMKN* αβ^−/−^ keratinocytes in comparison to the WT (Figure 3B). The protein abundance of dermokine and KRT10 decreased in both *DMKN* KO 3D cultures, confirming the results of the IHC staining (Figure 3a-b and Figure 2). KRT14, EGFR and ITGα6 showed no differential abundance in the *DMKN* βγ^−/−^ clone, as seen in the IHC analysis, but were slightly increased in the *DMKN* αβ^−/−^ clone (Figures 3A-3B and Figure 2). The epidermal proteases, KLK6 and aspartic peptidase retroviral like 1, known markers of pro-FLG processing, which itself regulates epidermal differentiation^9^, were significantly decreased in both *DMKN* KO 3D cultures (Figures 3A-B and Figure 2). Gene enrichment (GO) analysis of significantly decreased proteins from both the *DMKN* KOs compared to WT clones showed the enrichment of pathways such as keratinocyte differentiation and formation of the cornified envelope (Figure 3C), whereas the GO analysis of significantly increased proteins showed the enrichment of integrin binding, positive regulation of locomotion and growth factor binding pathways (Figure 3D).

**Figure 3:**
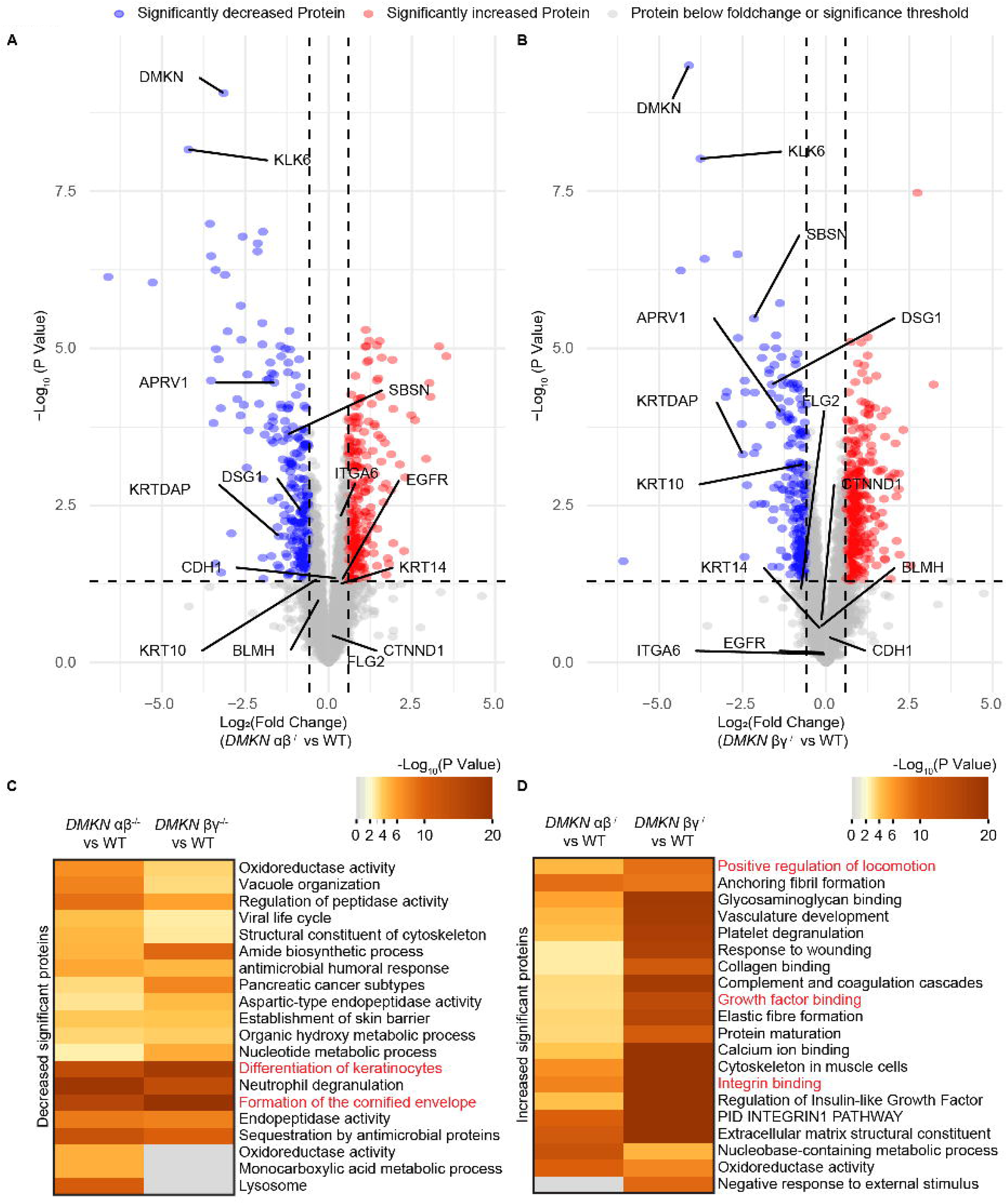
Proteomic analysis of dermokine-ablated 3D organotypic skin cultures shows phenotypic differences. **(A and B)** Volcano plots showing the differential protein expression between *DMKN* αβ^−/−^ **(A)** or *DMKN* βγ^−/−^ **(B)** and WT in 3D organotypic skin cultures. Known proteins commonly found in different epidermal strata are highlighted^1^. Values are Log_2_ transformed fold changes and -Log_10_ transformed p values (moderated contrast *t* test) of three different (3D) organotypic skin cultures. **(C and D)** Gene Ontology (GO) analysis of significantly decreased **(C)** or increased **(D)** proteins from *DMKN* αβ^−/−^ **(A)** or *DMKN* βγ^−/−^ **(B)** relative to the WT proteome.

Altogether, this data suggests that dermokine regulates the formation of the suprabasal but not the basal layer of epidermis without affecting early differentiation stages of keratinocytes through changes in protein abundances of epidermal differentiation markers and proteases based on our model of keratinocyte differentiation.

### Phosphoproteomics analysis reveals that dermokine regulates cell-cell adhesion proteins

To uncover how dermokine affected the observed phenotype of keratinocytes we studied changes in cellular signalling by deep quantitative proteomics and phosphoproteomics analyses of WT, *DMKN* αβ^−/−^, and *DMKN* βγ^−/−^ keratinocytes in regular 2D culture systems. (here referred to as endogenous *DMKN* αβ^−/−^ and *DMKN* βγ^−/−^, respectively)^14^. To focus on specific changes caused by the loss of dermokine, we also analysed the proteomes and phosphoproteomes of *DMKN* αβ^−/−^ or *DMKN* βγ^−/−^ keratinocytes treated with recombinant dermokine-β for 60min (here referred to as supplemented *DMKN* αβ^−/−^ and *DMKN* βγ^−/−^, respectively)^27^ (Figure S3A).

We firstly focused on the proteome and found 5118 unique quantified proteins with the median CV of the protein groups being below 20 % for all samples and the correlation analysis showing high correlation between replicates, whereas the conditions group separately well from each other in the PCA analysis (Figure S3B-S3D). Both the endogenous and the supplemented *DMKN* αβ^−/−^ or *DMKN* βγ^−/−^ proteomes differ from the endogenous WT proteome as shown by hierarchical clustering (Figure S4Aa). The similarity between the proteome of supplemented *DMKN* αβ^−/−^ and *DMKN* βγ^−/−^ and that of endogenous WT proteome is in line with previous findings where the proteome does not change after a short incubation time with an endogenous protein^28^. When we subjected the significantly differential proteins whose abundance decreased or increased in *DMKN* αβ^−/−^ and *DMKN* βγ^−/−^ to WT keratinocytes to GO enrichment analysis we found that the term “cadherin binding” was enriched in two conditions: when comparing the most abundant proteins from endogenous *DMKN* αβ^−/−^ and *DMKN* βγ^−/−^ to those from WT keratinocytes (Figure S4B) and when comparing the significantly decreased proteins from the supplemented *DMKN* αβ^−/−^ and *DMKN* βγ^−/−^ to those from endogenous keratinocytes (Figure S4C). These results suggest that proteins associated to “cadherin binding” are significantly more abundant in endogenous *DMKN* αβ^−/−^ and *DMKN* βγ^−/−^ keratinocytes compared to all the other conditions. This data also confirms that dermokine globally affects the keratinocyte proteome as shown for the 3D culture proteomes (Figure 3).

Next, we analysed the phosphoproteome of both endogenous and supplemented *DMKN* βγ^−/−^ and *DMKN* αβ^−/−^ as well as WT keratinocytes (Figure S3A) by enriching samples for phosphorylated peptides using automated Fe(III)-NTA, and a modified HRMS^1^-DIA strategy to cover a higher mass range (400-1400 m/z)^29–31^ (Figure S5A). In total, we identified 11778 phosphorylation sites after filtering for localization site probability (≥ 0.75) and 4453 fully quantifiable phosphorylated sites across all conditions on 2695 proteins (Figure S5B). All replicates showed high correlation scores (Figure S5C), and we found 10027 single, 1367 double and 384 multiple phosphorylated sites (Figure S5D) and 10315, 1164 and 299 phosphorylated serines, threonines and tyrosines, respectively (Figure S5E). Overall, the quality of the phosphoproteomics datasets was consistent with previous publications^12,31^. We observed significant differences between the phosphoproteome of endogenous *DMKN* βγ^−/−^ and *DMKN* αβ^−/−^ compared to the WT phosphoproteome (4453 phosphorylated sites) by hierarchical clustering (Figure S5F). The phosphoproteome of supplemented *DMKN* βγ^−/−^ and *DMKN* αβ^−/−^ keratinocytes was similar to the phosphoproteome of WT which suggests a considerable reversal of the phosphorylation pattern in *DMKN* βγ^−/−^ and *DMKN* αβ^−/−^ upon treatment with recombinant dermokine back to the WT phosphorylation pattern (Figure S5F).

To study how dermokine isoforms affected cellular signalling we applied a global Kinase-Substrate prediction approach, which reveals the likelihood of a kinase to modify a phosphorylated site by linking kinase recognition sequence motifs and known *in vivo* and *in vitro* phosphorylation profiles^32^, to the 4453 phosphorylated sites identified in the WT and the two KO samples. This approach identified 94 putative kinases whose downstream phosphorylated sites were visualized in a heat map (Figure 4A). Among the 94 kinases and based on the kinase activity score, we identified 30 kinases with a high change and 15 kinases which showed significantly highest changes between the endogenous WT and all the *DMKN* βγ^−/−^ and *DMKN* αβ^−/−^ samples (Figure 4b). Four of the latter kinases were tyrosine kinases (DYRK1A, PTK2B, SRC and SYK) of which both SRC and SYK showed decreased activity scores in the *DMKN* βγ^−/−^ and *DMKN* αβ^−/−^ keratinocytes (endogenous) compared to WT (Figure 4B). Next, we used the kinase-substrate annotations generated by the global kinase-substrate prediction approach to cluster substrates sharing similar kinase profiles and regulation^32^ and we identified 8 clusters (Figure S6). Interestingly, each clusters contained a different distribution of phosphorylated serine, threonine and tyrosines, sites ranging from cluster 2, 3, 4, and 6 not containing any phosphorylated tyrosine site, to cluster 8 containing only phosphorylated serine sites, and clusters 5 and 7 mainly containing phosphorylated tyrosine and threonine sites with cluster 7 being associated with PTK2B, SRC and SYK activity (Figure S6). GO enrichment analysis of the 8 clusters revealed a broad range of GO terms related to RNA regulation with the highest gene ratio in cluster 7, which, together with cluster 2, also contained several proteins associated with adherens junctions and cadherin binding (Figure 4C). This finding is in line with SRC, a known player in cell-cell adhesion^33^, being the most likely kinase regulating the proteins belonging to cluster 7 (Figure S7) and with cadherin binding being the term mostly dysregulated in the *DMKN* βγ^−/−^ and *DMKN* αβ^−/−^ keratinocytes compared to WT (Figure S4). To identify potential dermokine-related phosphorylated sites we counted the number of sites on each phosphorylated protein belonging to the pathways associated with cadherin binding and adherens junctions associated to clusters 2, 4, 5 and 7. We found several phosphorylated sites (between 8 and 12) on proteins associated to cell-cell adhesion, including 182 kDa tankyrase-1-binding protein (TNKS1BP1 (12)), tight junction protein 1 (TJP1 (11)), scribble (SCRIB (10)), tight junction protein 2 (TJP2 (10)), catenin β 1 (CTNNβ1 (9)), afadin (AFDN (8)) and catenin δ 1 (CTNNδ1 or p120 (8)) (Figure S6). Among these proteins we then focused on p120 as it was phosphorylated on S^252^, S^268^, S^288^ and T^916^, all playing crucial roles in modulating cell-cell adhesion complexes and cytoskeletal dynamics^20,34,35^ and due to the main GO cellular component and biological process terms associated with p120 being only “adherens junctions, cadherin binding and general cell adhesion”, as opposed to the multi-functional CTNNβ1^20,34,36–38^. To confirm that the regulated phosphorylated sites identified on p120 as well as other proteins were *bona-fide* phosphorylated sites due to changes in kinase activity and independent of translational regulation^39^ we normalised the phosphoproteome on the proteome (Figure S7A). We found that the normalised phosphoproteome of endogenous WT differed from that of endogenous *DMKN* βγ^−/−^ and *DMKN* αβ^−/−^ and that, when the *DMKN* βγ^−/−^ and *DMKN* αβ^−/−^ keratinocytes were supplemented with recombinant dermokine, their phosphoproteome reverted to WT keratinocytes (Figure S7A). This finding was consistent with what we observed for the phosphoproteome before normalization (Figures 5A-5B). Importantly, the phosphorylated p120 sites were still upregulated in supplemented *DMKN* βγ^−/−^ and *DMKN* αβ^−/−^ keratinocytes compared to respective endogenous KO samples (Figure S7b), confirming the phosphorylation of p120 downstream of dermokine.

**Figure 4:**
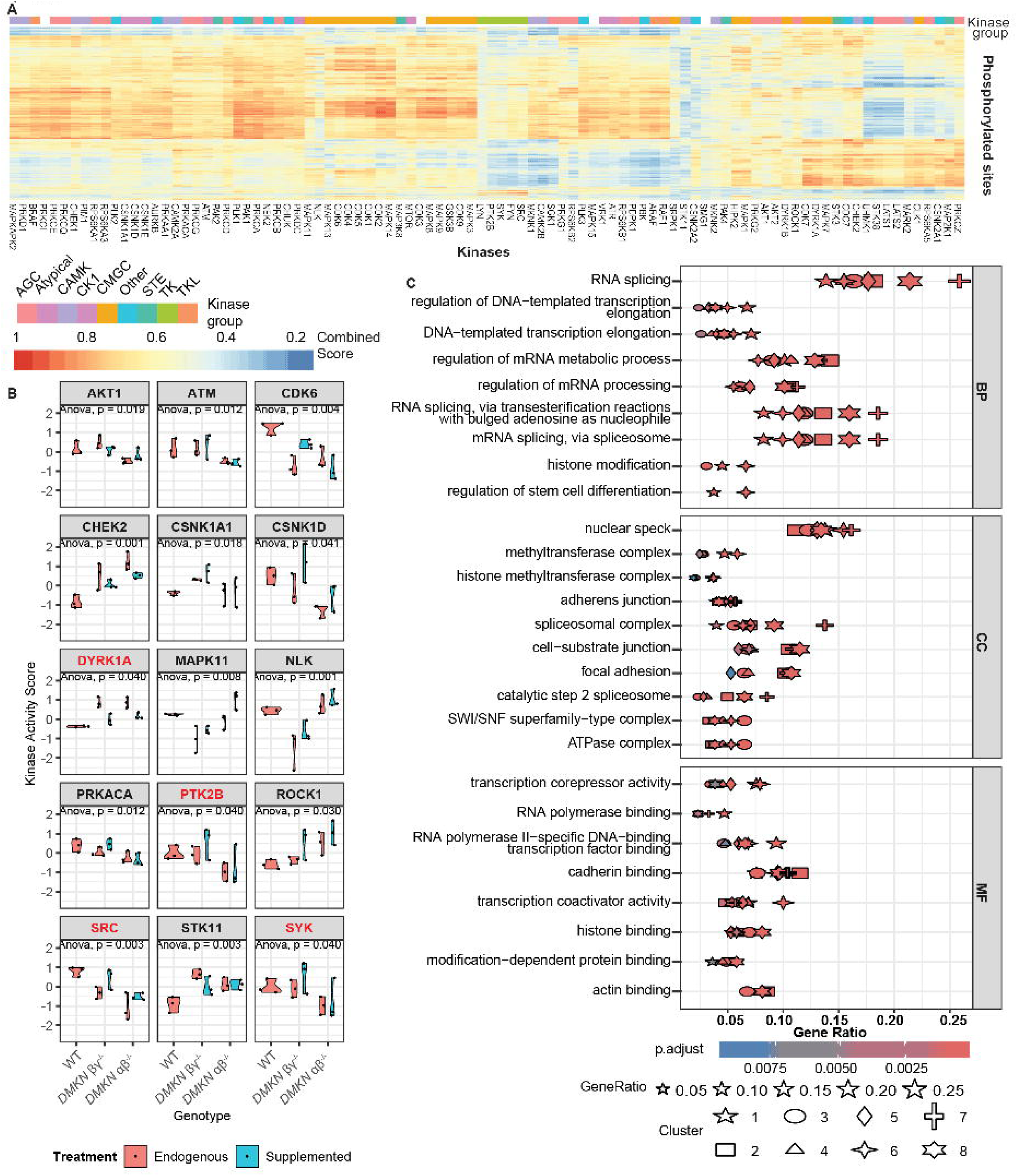
Absence of dermokine results in phosphorylation changes in adhesion proteins. **(A)** The Kinase-Substrate prediction approach^32^ matched quantified phosphorylated sites in the WT and the two KO samples with the 94 identified putative kinases. The heatmap shows the combined score of the *in vivo* or *in vitro* kinases and sequence recognition motifs^32^. Vertical axis shows substrates, and horizontal axis represents kinases. Family of kinases are color-coded on top of the graph. **(B)** The kinase activity score for each endogenous and supplemented *DMKN* αβ^−/−^, *DMKN* βγ^−/−^ and WT condition is shown for 15 significant kinases out of the 94 identified kinases. Values are kinase activity scores from N=3 technical replicates from the different genotypes and treatments. One-way ANOVA test determined changes among all genotypes. **(C)** Gene ontology analysis from proteins whose phosphorylated sites were annotated to kinases identified in clusters shown in Figure S6.

**Figure 5:**
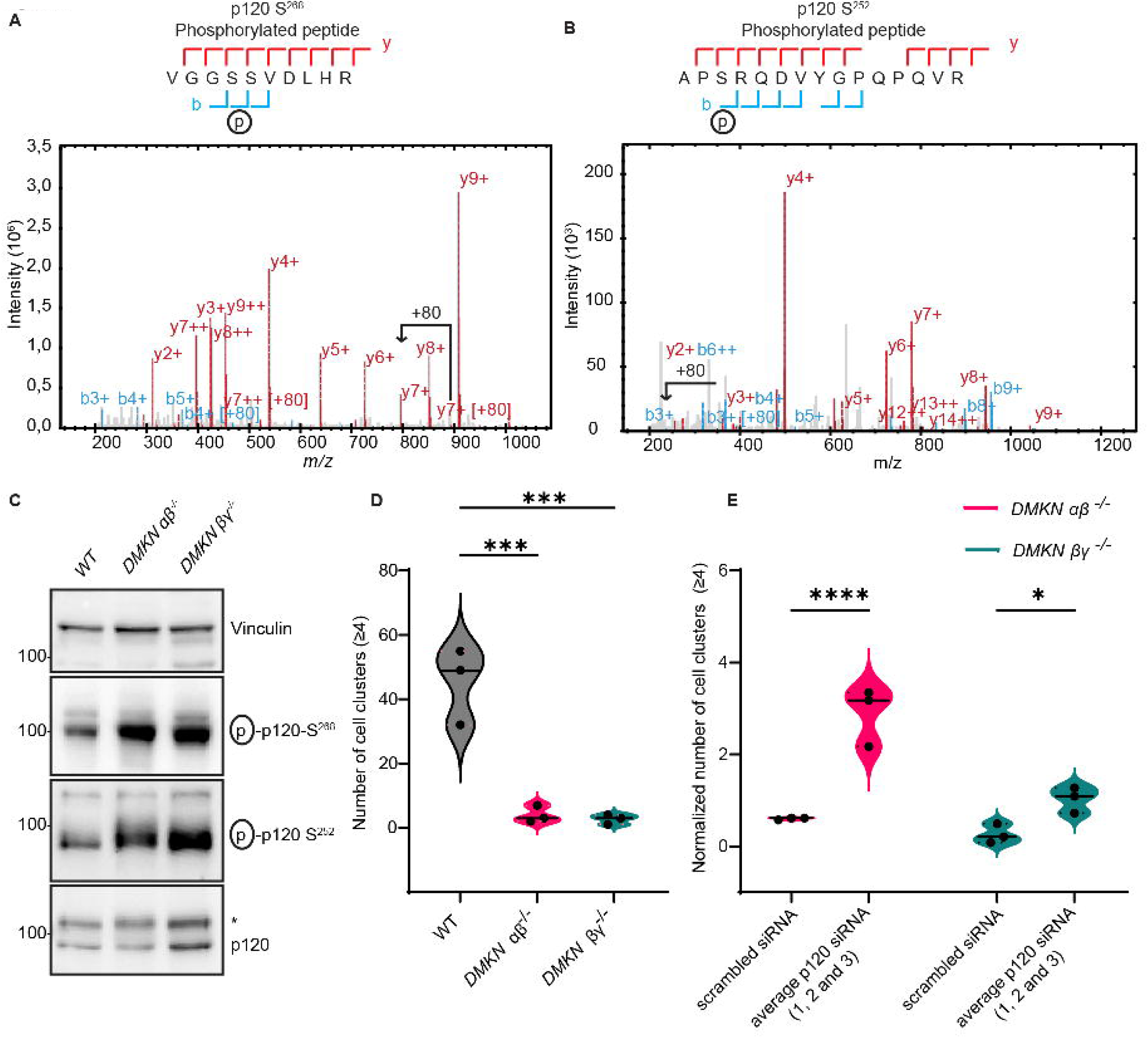
p120 phosphorylation increases and cell-cell adhesion decreases in dermokine*-*depleted keratinocytes. **(A and B)** MS/MS spectra of the tryptic phosphorylated peptide VGGpSSVDLHR from p120 at S^268^ **(A)** and APpSRQDDVYGPSQPQVR from p120 at S^252^ **(B)**. **(C)** Immunoblotting with the indicated antibodies of p120 phosphorylation (S^252^ and S^268^) in human *DMKN* αβ^−/−^, *DMKN* βγ^−/−^ and WT keratinocytes. **(D)** Quantification of 4 or more cell clusters from the cell-cell adhesion assay using either human *DMKN* αβ^−/−^, *DMKN* βγ^−/−^ and WT keratinocytes. Cell cluster (≥4) were counted, and each replicate was visualized as dot in the violin plot. *** = P < 0.001 (one-way ANOVA and a Fishers Least Significant Difference test). **(E)** *DMKN* αβ^−/−^, *DMKN* βγ^−/−^ and WT keratinocytes transfected for 72 h with p120 siRNA (1, 2 and 3) as well as scrambled siRNA were subjected to the cell-cell adhesion assay. The number of cell cluster (≥4) of *DMKN* αβ^−/−^ and *DMKN* βγ^−/−^ keratinocyte were counted and normalized to the average WT number of cell cluster within the same siRNA treatment. Shown are the siRNA averages for each genotype. Values are N = 3 biological replicates. * = P < 0.05, **** = P < 0.0001 (two-way ANOVA and a Fishers Least Significant Difference test).

In summary, proteomics and phosphoproteomics analyses identified key phosphorylated proteins such as p120 with a known role in cell-cell adhesion as being dysregulated downstream of dermokine.

### Dermokine decreases cell-cell adhesion through p120

To study the role of p120 downstream of dermokine, we checked the phosphorylation of p120 on two of the identified phosphorylated sites, serine 252 (S^252^) and serine 268 (S^268^), for which antibodies and fully annotated MS/MS spectra were available (Figures 4A-4B). Immunoblotting analysis of lysates from endogenous *DMKN* βγ^−/−^, *DMKN* αβ^−/−^ and WT human keratinocytes showed that the phosphorylation of both S^252^ and S^268^ on p120 increased in the absence of dermokine, whereas the total level of p120 was not affected by the absence of dermokine (Figure 5C). Furthermore, we observed that p120 phosphorylation levels remain similar to WT when recombinant dermokine was supplemented to *DMKN* βγ^−/−^, *DMKN* αβ^−/−^ and WT keratinocytes (Figure S8a). As p120 is known to regulate cell-cell adhesion^35^ and cell-cell adhesion was one of the dysregulated GO terms downstream of dermokine (Figures 4C, 5C-5E, S4A-S4B, Figure S7A-S7B) we tested whether dermokine regulated cell-cell adhesion in keratinocytes by a cell-cell adhesion assay^40^. We found that dermokine-ablated keratinocytes significantly form less cell clusters compared to WT keratinocytes (Figures 5D and S8B). Then, we tested whether p120 affected dermokine-mediated cell-cell adhesion in keratinocytes by performing the cell-cell adhesion assay in keratinocytes depleted of p120 using small interfering RNA (siRNA). We confirmed p120 knockdown in WT as well as in *DMKN* βγ^−/−^and *DMKN* αβ^−/−^ keratinocytes by immunoblotting (Figure S8C) and found that both *DMKN* αβ^−/−^ and *DMKN* βγ^−/−^ keratinocyte depleted of p120 significantly increased the number of cell cluster relative to p120-deficient WT keratinocytes (Figure 5E). These findings suggest that dermokine regulate cell-cell adhesion via p120 in keratinocytes.

As dermokine has been previously associated to skin disorders^41^, we stained skin biopsies from healthy and psoriatic skin, poorly- and well-differentiated SCCs as well as non-healing wounds, for dermokine as well as for p120 expression (Figures 6, Figure S9). We observed dermokine expression in healthy skin, psoriasis, poorly- and well-differentiated SCCs (Figures S9A-C) but the lowest dermokine expression in non-healing wounds (Figures 6A-B and Figure S9D). We quantified differential dermokine expression relative to the distance to the wound bed in healthy and non-healing wounds and found that non-healing wound-edge keratinocytes showed significantly lower dermokine expression across the epidermal profile, whereas, within the same biopsy, healthy keratinocytes showed no changing horizontal dermokine expression (Figure 6C, Figure S9E). To visualize where keratinocytes expressed dermokine, we fitted a non-linear regression curve previously used to explain differences in patient data from non-healing wounds and healthy skin^42^. We demonstrate that the average rate (K(Fast)) at which dermokine expression decreases, was higher in non-healing wound-edge keratinocytes (K(Fast) = 4.4E-03) compared to healthy skin keratinocytes (K(Fast) = 5.7E-08) (Figure 6C). Therefore, our data show less dermokine expression in wound-edge keratinocytes which suggests decreased cell-cell adhesion^7,8^ and confirms the phenotype of dermokine-depleted keratinocytes *in vitro* (Figure 5). Interestingly, p120 expression remained similar across all the analysed samples (Figure 5d and Figure S9f), as shown in keratinocytes *in vitro* (Figure 5c). To evaluate changes in phosphorylated p120 we used two antibodies specific for staining: one against phosphorylated S^252^ to recapitulate the immunoblotting findings (Figure 5C) and one against T^310^, as independent phosphorylated site on p120, also quantified in the phosphoproteomics dataset. Both these phosphorylated sites are known to inhibit cell-cell adhesion and induce cell motility^20,36,43^. After normalization of phosphorylated over total levels of p120 we found a significant increase of phosphorylated p120 on S^252^ and T^310^ in non-healing wounds compared to healthy skin (Figures 6E-F), confirming a link between dermokine, p120, and cell-cell adhesion.

**Figure 6:**
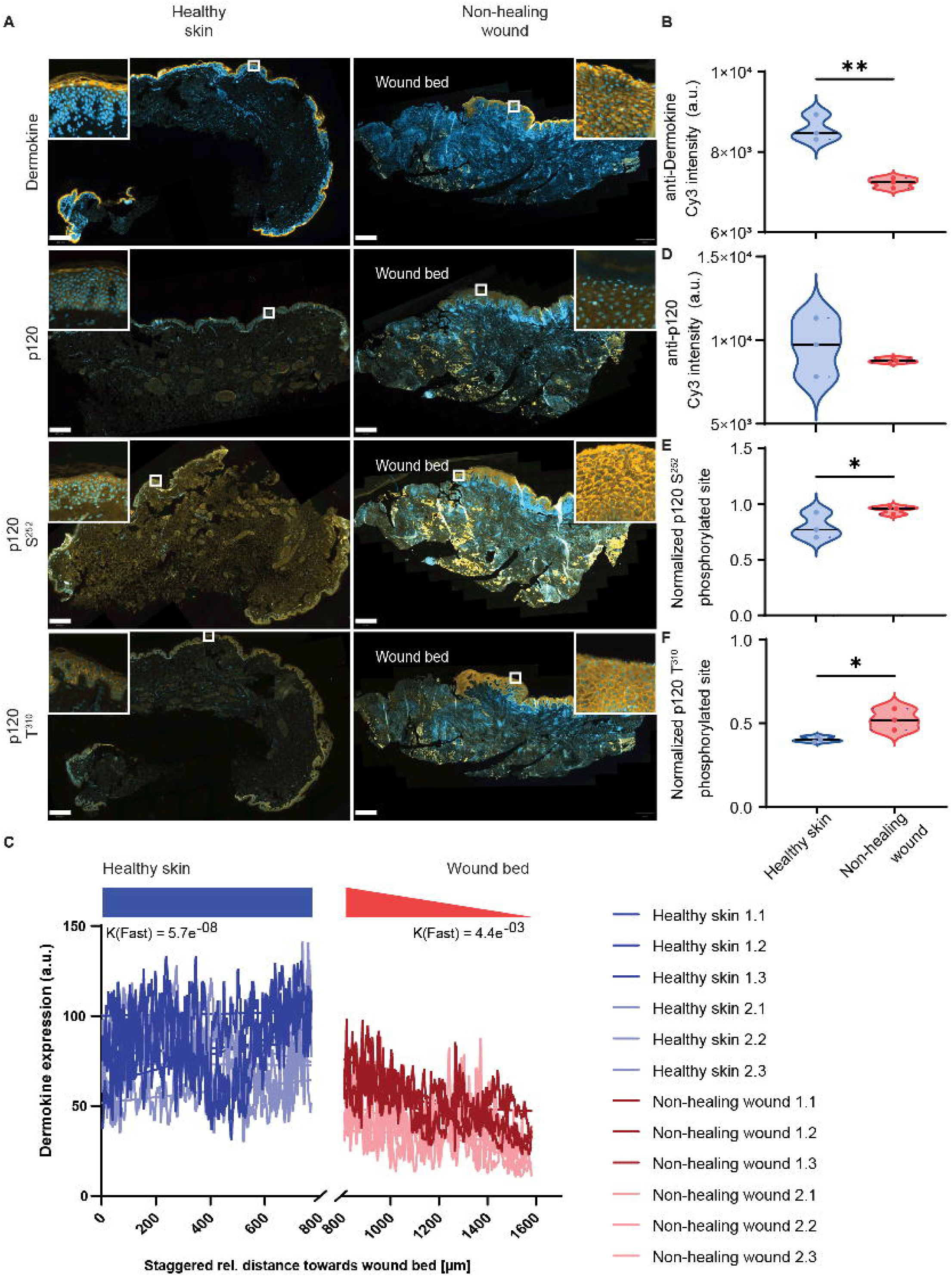
Dermokine expression decreases and p120 phosphorylation increases in non-healing wounds. **(A)** Representative images of immunofluorescent staining from biopsies of healthy skin and non-healing wounds from venous leg ulcers. (**B, C, E and F)** Quantification of immunofluorescent staining with the indicated antibodies from **(A)**. Values are the median Cy3 intensity values of N = 3 samples from different anatomical locations (left and right leg) of the same patient (non-healing wounds) and of three patients (healthy skin). Scale bar for the healthy skin is 400 µm and for non-healing wounds 1 mm. The scale bar for the magnified images is reported in extended figure 1. * = P < 0.05, ** = P < 0.01 (one-tailed, unpaired *t* test). **(D)** Quantification of dermokine intensity across 770,25 µm from the wound-edge towards healthy keratinocytes. The quantified area was measured in three horizontal rectangles across the epidermal profile (Figure S9D). Values are Cy3 anti-dermokine intensities along 3 equally shaped rectangles placed within the same biological sample at two different anatomical sites.

In conclusion, multi-omics analyses of dermokine-depleted human keratinocytes showed that the absence of dermokine isoforms increases phosphorylated p120 and inhibits cell-cell adhesion *in vitro* (Figure 5). Dermokine-dependent regulation of cell-cell adhesion was also observed in keratinocytes derived from healthy and diseased patients which showed decreasing dermokine expression and increased phosphorylation of p120 in non-healing wounds compared to healthy samples (Figure 6). As increased abundance of these phosphorylated sites has been reported to inhibit cell-cell adhesion and increase cell motility^20,34^, we suggest a link between lowered dermokine expression, increased p120 phosphorylation, and cell-cell adhesion regulation in non-healing wounds (Figure 7).

**Figure 7:**
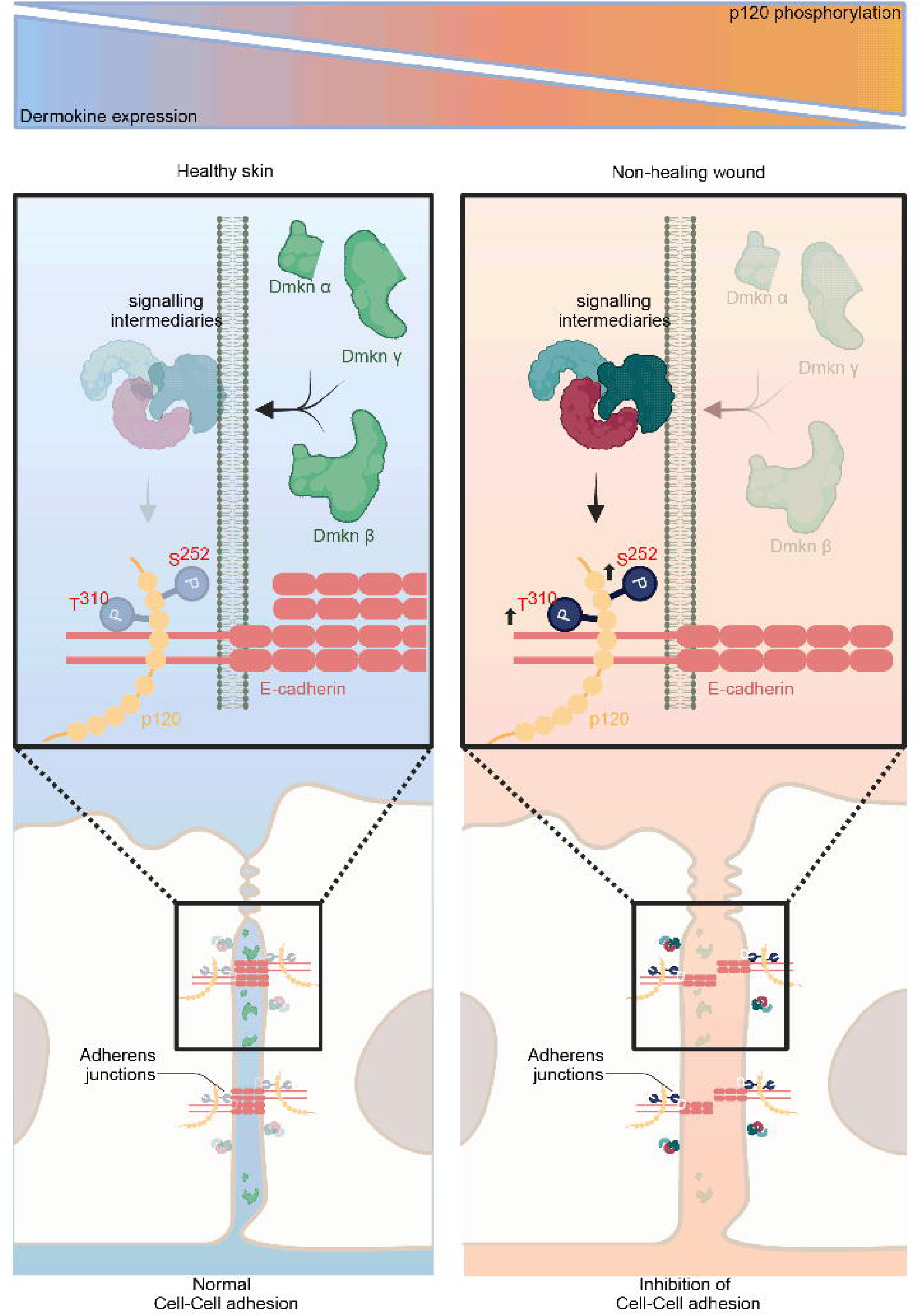
Model of dermokine function in human keratinocytes. Relationship between dermokine and cell-cell adhesion relative to p120 phosphorylation, illustrated as gradient. Depletion of dermokine leads to high p120 phosphorylation levels inhibiting p120 binding to the juxtamembrane domain of E-Cadherin, preventing cell-cell adhesion. This Figure was created in Biorender.

## Discussion

This study combines genome engineering, multi proteomics analyses of 3D organotypic cultures, and staining of patient derived samples to elucidate the functions of dermokine in keratinocytes. The MS-based proteome, targeted proteome, and phosphoproteome datasets provide insights into the function of dermokine in both regular 2D and 3D skin organotypic culture derived from *DMKN* αβ^−/−^, *DMKN* βγ^−/−^, and WT keratinocytes. In contrast to earlier studies, which relied on mRNA quantification to differentiate between isoforms^14^, we were able to quantify domain-specific proteotypic peptides via targeted proteomics and suggest a robust alternative to immunoblotting for validation of protein ablations^44^. Subsequently, we used organotypic cultures to investigate the effects of isoform specific loss of dermokine in keratinocytes in epidermal architecture. The overall thinner epidermis-like structure in *DMKN* αβ^−/−^ and *DMKN* βγ^−/−^ 3D cultures indicate aberrant keratinocyte differentiation ^41,45^. Our data support previous evidence^14^ that dermokine-isoforms indirectly regulate the expression of suprabasal proteases and TGMs, responsible for keratinocyte differentiation^9^. Interestingly, our data revealed no direct difference between dermokine isoforms in human keratinocytes. However, we discovered a functional overlap of the three main dermokine isoforms, showing no clear evidence that residual dermokine-α compensates for the loss of dermokine-β, as described in mice^14^.

To uncover the function of dermokine in keratinocytes, we acquired the phosphoproteome of *DMKN* αβ^−/−^, *DMKN* βγ^−/−^ and WT keratinocytes, by optimized DIA assays, based on a semi-automated phosphorylated peptide enrichment, which allowed us to substantially reduce our input material, from commonly used 3-5 mg to 200 µg lysate^31^. Despite the relatively low amount of starting material, we could fully quantify almost 5000 phosphorylated sites. The semi-automated sample preparation and increased mass range acquisition strategy has the potential to benefit other research groups, who might be limited by the amount of biological material, like FFPE tissues or single cells^46,47^. In this study, we identified p120 phosphorylation, associated with cadherin binding and adherens junctions, downstream of dermokine. Proteomic analysis corroborated increased levels of cadherin-binding proteins downstream of dermokine in keratinocytes. Through cell-cell adhesion assays, we showed a loss of cell cluster formation in *DMKN* αβ^−/−^ and *DMKN* βγ^−/−^ keratinocytes. Interestingly, by depleting p120 expression we reversed cluster formation to WT levels in *DMKN* αβ^−/−^ and *DMKN* βγ^−/−^ keratinocytes. Changes in cell-cell adhesion may be due either to p120 phosphorylation or to a lack of p120 expression, which could not be determined based on the transient interference of p120 expression. However, we stained phosphorylated p120 in healthy and diseased patient samples and correlated increased phosphorylated p120 with inhibition of cell-cell adhesion which may indicate a direct role of p120 phosphorylation in dermokine-dependent regulation of cell-cell adhesion both *in vitro* and *in vivo*^20,34,35^. A possible underlying mechanism might be that the p120-armadillo-repeat region, shortly after phosphorylated T^310^, may bind to the E-Cadherin juxtamembrane domain^20^. Conceivably, the increased p120 phosphorylation in the absence of dermokine may inhibit the binding of p120 to E-Cadherin by steric hindrances and induce protein turnover by internalization of E-Cadherin via endocytosis^35^. Our proteomics data did not show different levels of E-Cadherin, most likely due to the total cellular protein quantities remaining the same, regardless of if the protein is internalized or embedded in the membrane^34^. Our study further suggests that the kinases SRC, casein kinase 1 epsilon (CSNK1E), and ERK1/ERK2, which are known to phosphorylate p120 at Y^228^, S^268^, T^310^, respectively^20,34,48,49^ may be responsible for the dermokine-p120-E-cadherin-dependent phenotype.

Previous research investigating dermokine expression in human skin disorders has utilized an in-house anti-dermokine-βγ antibody which has only provided qualitative information^41^. Here, we used a commercially available anti-dermokine-β, and putative dermokine-γ antibody^50^ and quantified the stained area from healthy and diseased patient samples. Collectively, we found consistent dermokine expression in healthy, psoriatic skin and all SCCs, while non-healing wounds showed the highest dermokine expression changes. Contrary to our findings, Hasegawa *et al.* showed diminishing dermokine expression in poorly-differentiated SCCs and more dermokine expression in both psoriatic and wound tissue^41^. A possible explanation for the dramatic changes in wounds might be that we stained non-healing human wounds from venous leg ulcers, while Hasegawa *et al.* used a murine wound healing model^41^. This discrepancy supports the hypothesis that murine models may differ from human conditions, in part due to different immune cells repertoire and tissue architecture which may alter the healing process^51,52^. Another possible explanation for this discrepancy may be that non-healing wounds are chronically trapped in between the inflammatory and proliferative phase^53^. We show that keratinocytes at the non-healing wound edge fail to switch into either migratory or differentiating phenotype^6^. Our data improves our understanding of wound-edge keratinocytes, which in a regulated way undergo partially reversible EMT for re-epithelization, expanding the hypothesis that dysregulated wounds resemble cancer^54^. An important aspect to be considered here is that the proteome and probably the phosphoproteome composition of non-healing wounds most likely changes relative to healing wounds^55^. Conclusively, while dermokine expression decreased in non-healing wounds, phosphorylated p120 increased relative to healthy skin, which is consistent with previous studies reporting that cell-cell adhesion proteins, in venous leg ulcers, are mostly downregulated^6^.

Taken together, the functional multi-omics approach described here has the potential to improve our understanding of not only dermokine but also differential phosphorylation pattern in non-healing wounds. We could infer that non-healing wounds show low dermokine expression and high p120 phosphorylation, associating dermokine expression with cell-cell adhesion and potentially cell migration. In a broader perspective, our approach might guide drug development for cutaneous biology when integrating “-omics” techniques with functional assays^56^.

## STAR Methods

### Key resources table

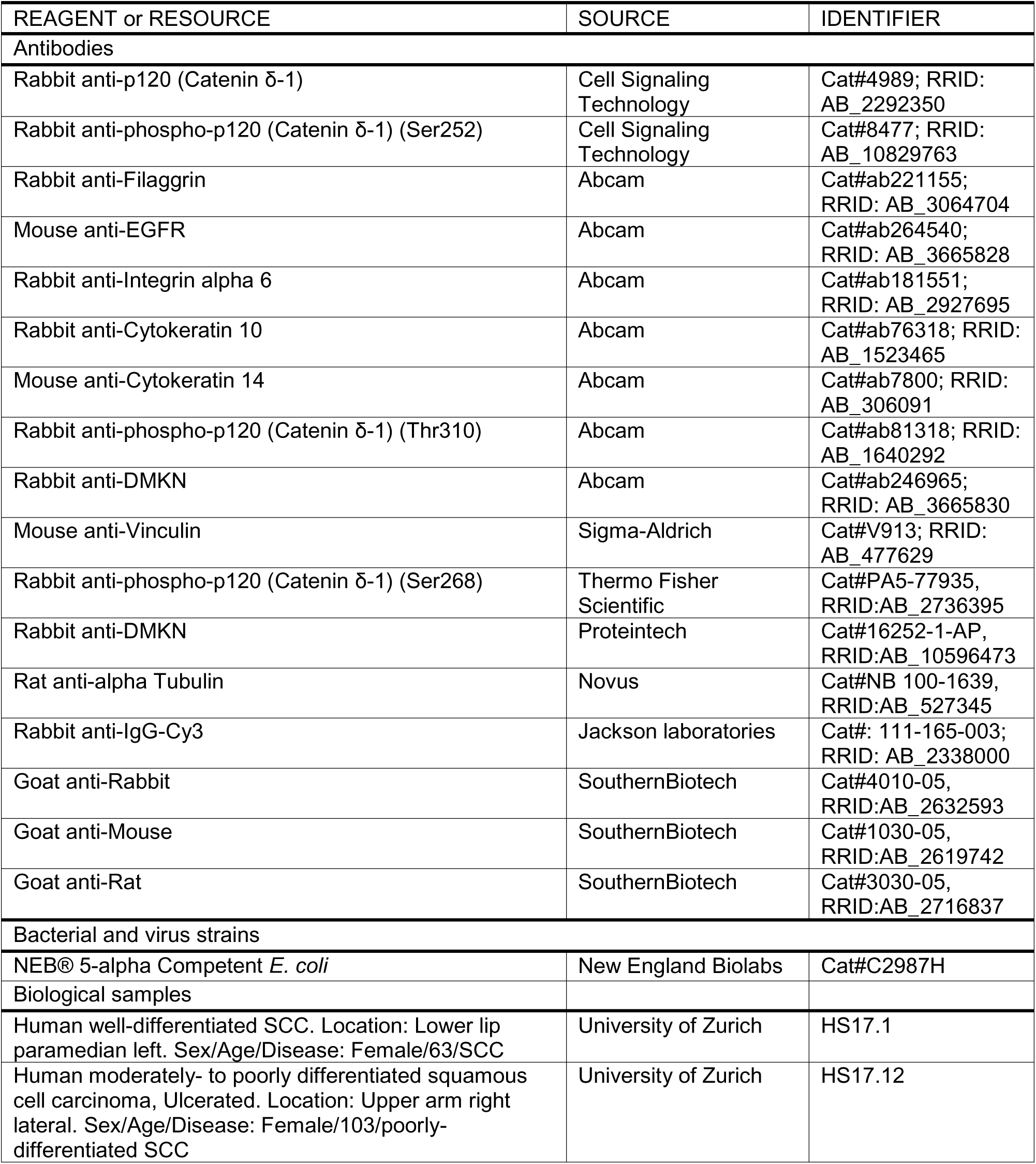
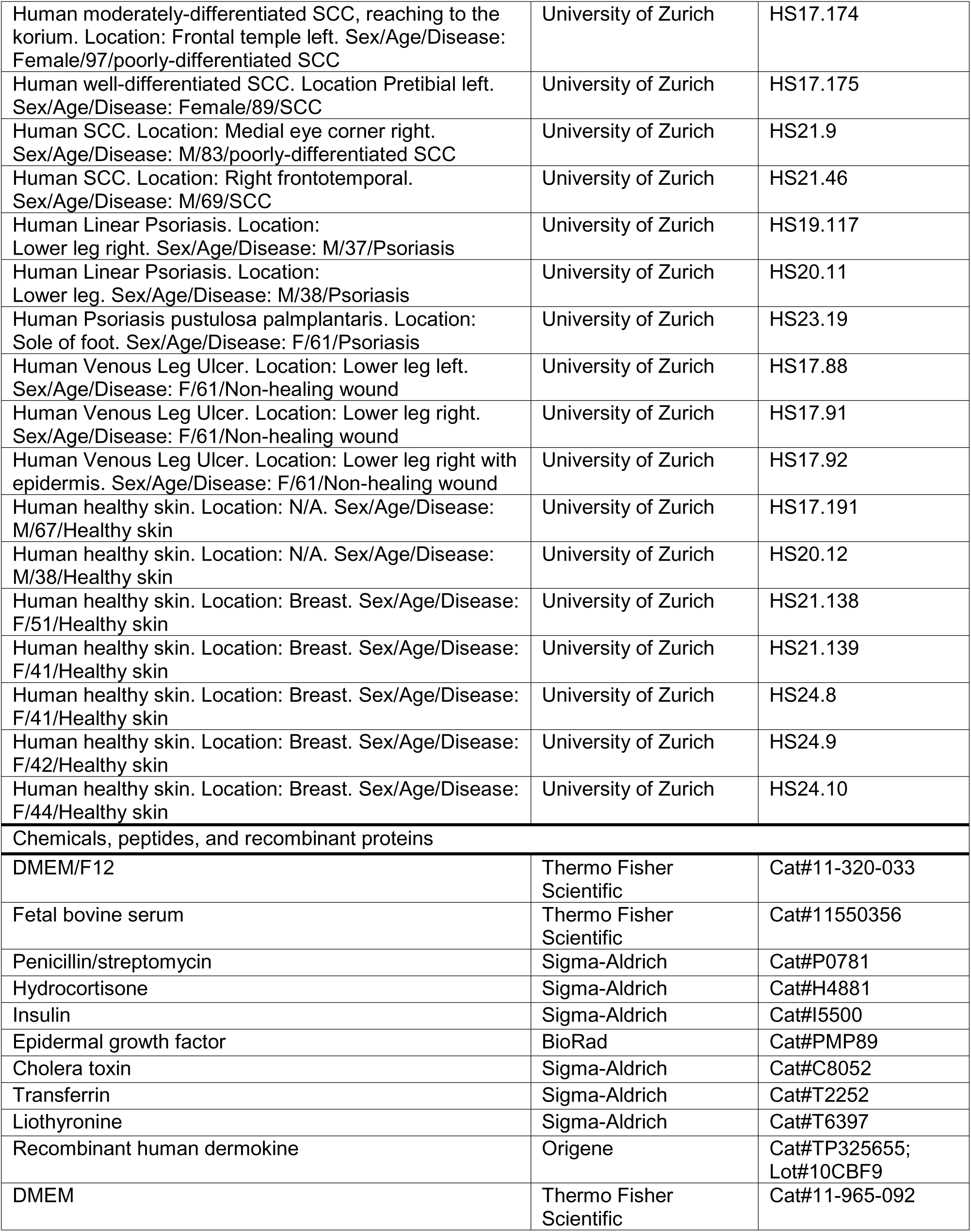
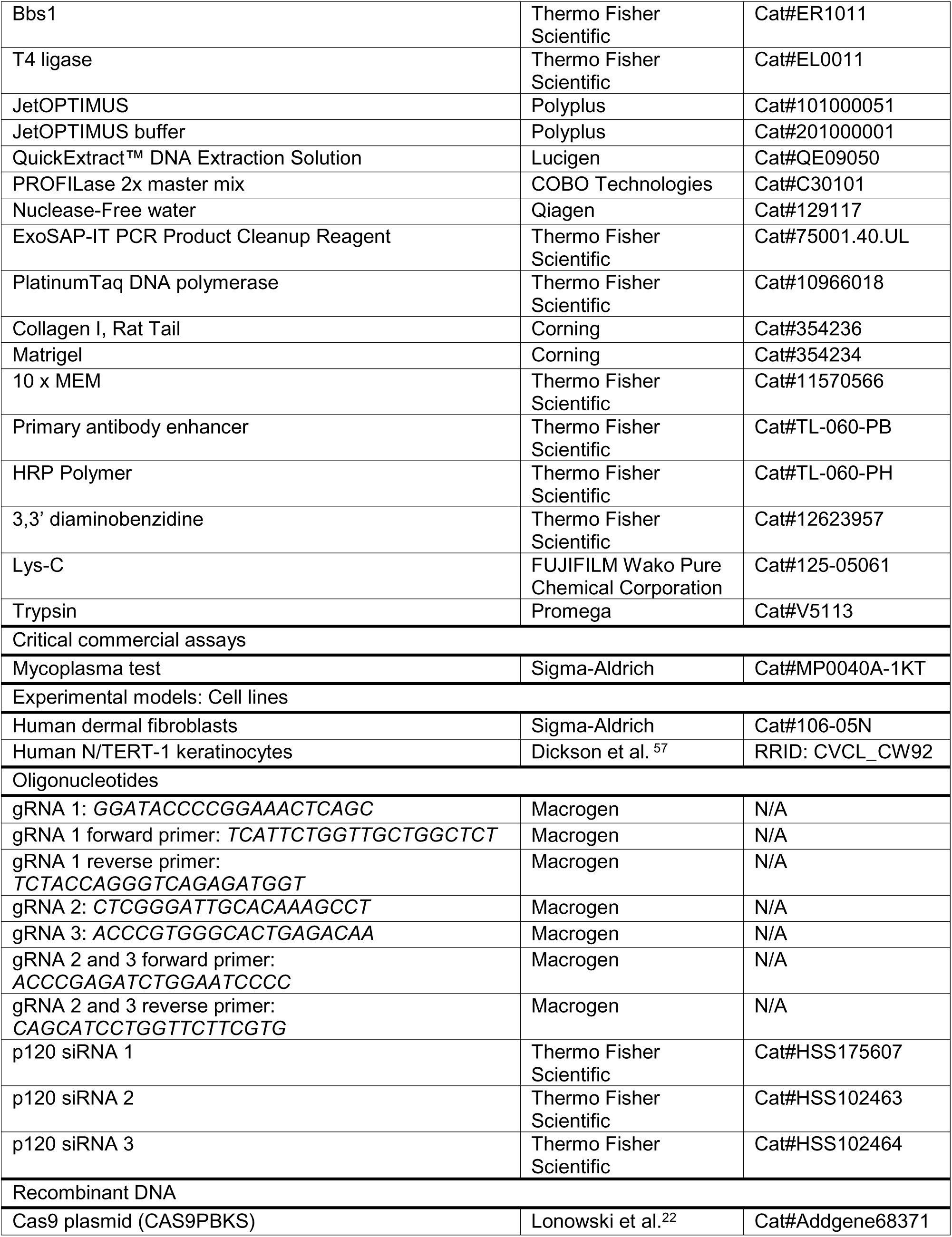
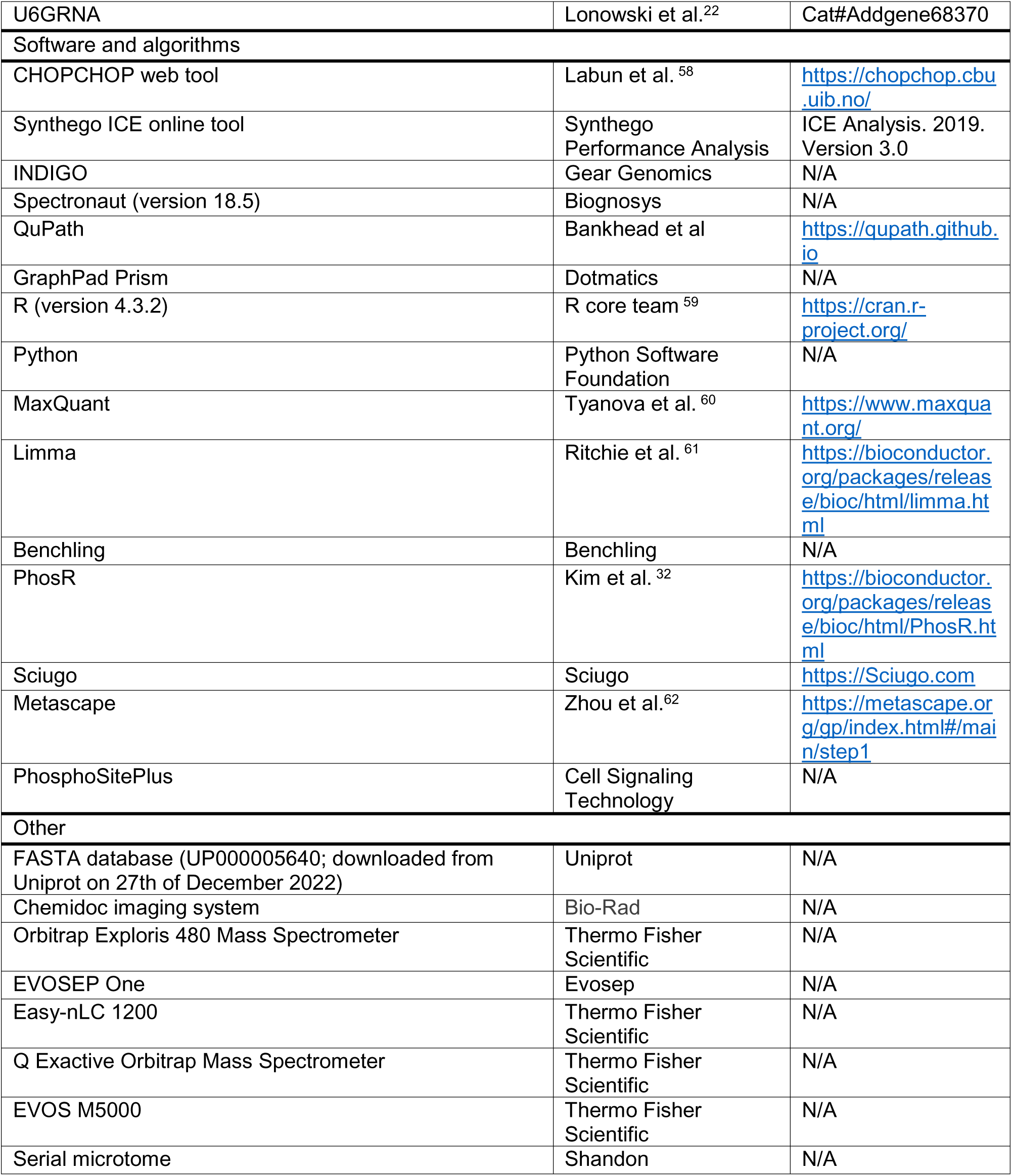

### Resource availability

#### Contact for Reagent and Resource Sharing

Further information and requests for resources and reagents should be directed to and will be fulfilled by the Lead Contact, Chiara Francavilla.

#### Materials availability

All data reported in this paper will be shared by the lead contact upon request.

#### Data and code availability

- Any additional information required to reanalyze the data reported in this paper is available from the lead contact upon request.

### Experimental Model and Subject Details

#### Cell culture

Human keratinocyte telomerase reverse transcriptase immortalized (h/TERT immortalized) N/TERT-1 cells derived from clinically normal foreskin tissue. N/TERT-1 keratinocytes were grown in DMEM/F12 growth medium supplemented with RM+ (DMEM/F-12 (Thermo Fisher Scientific), 10% fetal bovine serum (FBS) (Thermo Fisher Scientific), 1% penicillin/streptomycin (P/S) (Sigma-Aldrich), 0.4 µg/mL hydrocortisone (Sigma-Aldrich), 0.5 µg/mL insulin (Sigma-Aldrich), 10 ng/mL epidermal growth factor (BioRad), 0.1 nM cholera toxin (Sigma-Aldrich), 5 µg/mL transferrin (Sigma-Aldrich), 20 pM liothyronine (Sigma-Aldrich)) in a humidified incubator at 37 °C and 5% CO_2_. Human dermal fibroblasts were purchased from Sigma-Aldrich (Sigma-Aldrich) and grown in DMEM (Thermo Fisher Scientific), 10% FBS (Thermo Fisher Scientific) and 1% P/S (Sigma-Aldrich). All cell lines were tested mycoplasma negative before transfection using a commercial kit (Sigma-Aldrich).

#### Antibodies

The anti-dermokine antibody recognizes the last C-terminal 200 amino acids (AA) of dermokine-β^50^. The C-terminal 66 AA of dermokine-α are equivalent to the C-terminus of dermokine-β, while dermokine-α has 24 differing AA.

#### Guide RNA design

Guide RNAs (gRNAs) were designed utilizing the general settings of the CHOPCHOP web tool to target specific sequences associated with the isoforms of the dermokine gene.

For the chemical-transfection approach we used gRNA 1, targeting exon 2. For the Atomic-Force-Microscopy (AFM) based approach we used a multiplexed Cas9 strategy to delete exon 17.

#### Chemical transfection to generate *DMKN* βγ^−/−^ keratinocytes

We employed a two-plasmid system to enable the assembly of gRNA and Cas9 protein inside the cells for gene editing. The Cas9 plasmid (CAS9PBKS) was obtained from Addgene, while the gRNA plasmid was reconstructed using the U6GRNA (Addgene) vector for the expression of green fluorescent protein (GFP) and dermokine gRNAs (Macrogen). To insert the gRNA target we firstly used *Bbs1* (Thermo Fisher Scientific) to linearize the U6GRNA (Addgene) plasmid and then T4 ligase (Thermo Fisher Scientific). After selecting transformed *E. Coli* clones on ampicillin agar plates, successful insertion of the gRNA target was confirmed through Sanger sequencing (Macrogen). Keratinocytes were seeded at a density of 5e^5^ cells/mL in a 6 well-plate one day prior, to achieve 50-80% confluency on the day of transfection. Each transfection required 2 µg of CAS9PBKS. To use JetOPTIMUS® (Polyplus), 2 µg of CAS9PBKS were diluted in 200 µL JetOPTIMUS® buffer and 2 µL of JetOPTIMUS® reagent was added. The mixture was stirred well and incubated at room temperature for 10 min. Two days after transfection, we used the MA900 cell sorter (Sony Biotechnology) to isolate the top 10-15% of cells expressing GFP, which were subsequently cultured for up to 2 weeks to allow for cell expansion. Finally, single-cell sorting of GFP negative cells was performed using 96-well plates (Thermo Fisher Scientific). Genotyping was performed using PCR amplification and Sanger sequencing. The QuickExtract™ DNA Extraction Solution (LGC Biosearch Technologies) was utilized to extract DNA, which served as the template for PCR amplification. Approximately 1e^4^ cells were collected, washed with PBS, and resuspended in 30-50 µL of QuickExtract™ (LGC Biosearch Technologies) DNA Extraction Solution. The samples were heated at 70 °C for 20 min to extract DNA from cells, followed by 95 °C for 10 min. The PCR setup included 12.5 µL of PROFILase™ 2x Master Mix (COBO Technologies), 10 µL of Nuclease-Free water (QIAGEN), 0.25 µL of 25 µM forward primer, 0.25 µL of 25 µM reverse primer, and 2 µL of DNA template. The PCR tube was placed in an ABI 9902 Veriti™ thermocycler (Thermo Fisher Scientific). The PCR program consisted of three stages: Stage 1, the samples were heated to denature the double-stranded DNA. In stage 2, the samples underwent cycles of denaturation, annealing and extension. Stage 2 was repeated 38 times. In stage 3, the sample was heated to ensure complete synthesis of all ongoing templates. The PCR product was purified and sent to Macrogen for Sanger sequencing (Macrogen). The sequencing data were analysed using the Synthego ICE tool (Synthego Performance Analysis). Mutant clones were selected based on the presence of indels.

#### FluidFM™ based nano-injection to generate *DMKN* αβ^−/−^ keratinocytes

The principle of FluidFM™ is described in detail in^21^. A previous study used FluidFM™ to perform multiplex editing in CHO-K1 cells^63^. In brief, single keratinocytes were seeded into the centre of a well of a 12-well plate using a DispenCell™ (SEED Biosciences). The cells were left to adhere overnight and 50-100 fL of RNP solution was injected into the nucleus of single cells using FluidFM™ technology. The injection mixture consisted of 0.5 pmol gRNA (IDT), 0.366 pmol of Cas9 *(IDT)* and 20 ng of GFP mRNA (Milteny) per µL. Injection success was monitored 24 h after injection by fluorescent microscopy (GFP+ cells). Single cells were subsequently expanded using the medium described above. To improve single cell outgrowth, untreated keratinocytes were seeded into co-culture inserts and placed into the 12-well plates containing the injected cells. When cell colonies reached 1e^3^ cells, cells were detached and seeded into a 12-well plate. 48 to 72 h later, cells were split at a 1:3 ratio and a portion was used for isolation of genomic DNA. Genomic DNA was isolated from the clones using QuickExtract (Lucigen) according to the manufacturer’s instructions. PCR was performed using the PlatinumTaq DNA polymerase with GC enhancer (Thermo Fisher Scientific) and an α cycler (PCRmax) using 1 µL of DNA lysate. Primer concentrations were 0.4 µM each. For separation and visualization of PCR amplification products using 1.5% (w/v) agarose gels. Gels were casted and run using a cleaver scientific agarose gel system (Cleaver Scientific). DNA and size marker (4 µL, GeneRuler 100 bp) were separated at a constant voltage of 100 V for 25 min. Sanger sequencing was performed as follows: 7.5 µL of PCR product (50-300 ng) was purified using ExoSAPIT™ (Thermo Fisher Scientific) according to the instructions. The volume of the PCR product was adjusted to 12 µL with ultra-pure water and 3 µL of sequencing primer (20 µM) added to the reaction. Sanger sequencing was performed by Microsynth™ (Microsynth). Editing outcomes were determined by aligning the sequence trace files derived from potentially edited clones to the target sequence obtained from ENSEMBLE using the molecular biology suite of Benchling™ (Benchling). The sequence trace peaks surrounding the gRNA cut site were inspected visually and in case of overlapping Sanger peaks or visible deletions, the sequencing files were further analysed by sequence trace file deconvolution using INDIGO™ (Gear Genomics). From the deconvolved sequences, the individual alleles were reconstructed and the effects of the deletions/insertions on the reading frame of the targeted sequence were determined. Clones carrying frame shift mutations within the coding sequence of all detected alleles or deletion of the targeted exons were considered as knockouts (Figure S1).

#### Generation of human *DMKN* αβ^−/−^ and *DMKN* βγ^−/−^ keratinocyte 3D organotypic skin cultures

3D organotypic skin cultures were generated by placing 600 µL of collagen/Matrigel (Corning) matrix containing 1e^5^ human dermal fibroblasts (Sigma-Aldrich) (210 µL of collagen I (Corning), 210 µL of Matrigel (Corning), 59 µL of 10x MEM (Thermo Fisher Scientific), 59 µL of FBS (Thermo Fisher Scientific), and 59 µL of primary human fibroblasts (Sigma-Aldrich)) into a 12-well plate Transwell insert (Sigma-Aldrich) and incubating for 1 h at 37 °C. After polymerization of the dermis-like layer, 59 µL of either wildtype or *DMKN* αβ^−/−^ or *DMKN* βγ^−/−^ keratinocytes (containing 1e^6^ cells) were seeded on top of the matrix and RM+ growth media was added below the insert. After overnight incubation at 37 °C, the 3D cultures were airlifted and left to grow at an air-liquid interface for 15 days with daily media changes. After 15 days, the 3D cultures were cut in half and either fixed in 4% paraformaldehyde (PFA) (Sigma-Aldrich) for 30 min at RT and embedded in paraffin for IHC analysis or the other half was snap-frozen in liquid nitrogen for proteomics analysis.

#### Immunohistochemical staining of WT, *DMKN* αβ^−/−^ and *DMKN* βγ^−/−^ 3D organotypic skin cultures

Generation of sections and stainings were performed at the DTU Healthtech. Paraffin sections were cut with a serial microtome (4 µm) (Shandon) and dried overnight at 35 °C. IHC staining was performed according to the manufacturer’s instructions (Thermo Fisher Scientific). In brief, sections were heated at 60 °C for 60 min before deparaffination. The sections were deparaffinized, rehydrated and blocked with Ultra V Block (Thermo Fisher Scientific) for 5 min. Slides treated with anti-dermokine antibody (1:100 dilution) (Abcam) were directly incubated for 1 h and rinsed twice with tris buffered saline (Sigma-Aldrich) for 5 min. Slides stained with either Anti-FLG (1:100), anti-EGFR (1:100) or anti-ITGα6 (1:500) antibodies were pre-treated in EDTA buffer (10 mM Tris and 1 mM EDTA, pH 9.0). Slides stained with either anti-KRT10 (1:1000) or anti-KRT14 (1:500) antibodies were pre-treated in citrate buffer (100 mM citric acid and 100 mM sodium-citrate, pH 6.0). In detail, slides were boiled in either EDTA or citrate buffer for 15 min, cooled for another 15 min and washed twice with washing buffer before incubation with respective antibodies. Primary antibody enhancer (Thermo Fisher Scientific) was applied and incubated for 10 min before being washed 3 times with washing buffer for 5 min. HRP Polymer (Thermo Fisher Scientific) was applied and incubated for 15 min before being washed 3 times with washing buffer for 5 min. Slides were incubated with 3,3’ diaminobenzidine (DAB) (Thermo Fisher Scientific) for 10 min and washed in washing buffer for 5 min prior to rinsing under tap water. Hematoxylin (Sigma-Aldrich) was applied onto the slides for 15 seconds and rinsed under tap water for 5 min before adding Pertex mounting medium (Histoline). The image slides were captured using the EVOS M5000 imaging system in phase contrast mode, 20x magnification (Thermo Fisher Scientific).

#### Mass spectrometry (MS)-based proteomics

Lysis buffer (4M guanidine hydrochloride (GuHCl) and 250 mM N-2-hydroxyethylpiperazine-N-2-ethane sulfonic acid (HEPES) (pH 7.8)) was mixed with 3D organotypic skin cultures, homogenized using the steel beads (QIAGEN) and TissueLyser II (QIAGEN). The lysate was processed twice for 1 min each at varying frequencies (3 Hz to 30 Hz), beads were removed using a magnetic rack and the suspension was heated for 5 min at 95 °C. Following this, the samples underwent five cycles of sonication and cooling using the TissueLyser ii (QIAGEN). After centrifugation, the protein content was determined using Nanodrop (Thermo Fisher Scientific) and Bradford assays (Bio-Rad). The samples were reduced and alkylated on aliquots of 50 µg protein using 5 mM tris(2-carboxyethyl)phosphin (TCEP) and 20 mM 2-chloroacetamide (CAA). The samples were diluted with 50 mM HEPES buffer (pH 7.8) and incubated with Lys-C (FUJIFILM) for 4 h, 37 °C followed by trypsin (Promega) for 16 h at 37 °C. The enzyme activity was blocked by 1% tri-fluoroacetic acid (TFA) and samples were desalted using Solaµ plates (Thermo Fisher Scientific), according to manufacturer’s instructions.

For each resuspended proteome, peptides were analysed using the pre-set ‘30 samples per day’ method on the EvoSep One instrument. Peptides were eluted over an EvoSep One defined 44-min gradient and analysed by mass spectrometry, Orbitrap Exploris 480 (Thermo Fisher Scientific), adapted from with the following settings: Spray voltage was set to 2.3 kV, funnel RF level at 40, and heated capillary at 240 °C. Full MS spectra were collected at a resolution of 120000, with an AGC target of 300% or maximum injection time set to ‘custom’ and a scan range of 350–1650 m/z. The MS^2^ spectra were obtained in DIA mode in the Orbitrap operating at a resolution of 15000, equipped with FAIMS Pro™ Interface (Thermo Fisher Scientific) with a CV of −35, −55 and −75 V, with an AGC target 1000% or maximum injection time set to ‘auto’, a normalised HCD collision energy of 30. The isolation window was set to 38 m/z with a 1 m/z overlap and window placement off. MS performance was verified for consistency by running complex cell lysate quality control standards, and chromatography was monitored to check for reproducibility.

Raw files were analysed using Spectronaut™ (Biognosys) directDIA+ (deep) default settings, spectra were matched against the homo sapiens reference FASTA database (UP000005640; downloaded from Uniprot on 27th of December 2022). Default settings were used without imputation. Dynamic modifications were set as Oxidation (M), Glutamine to pyro-Glutamine and Acetyl on protein N-termini. Cysteine carbamidomethyl was set as a static modification. Protein quantitation was done on the MS^2^ level, data filtering set to default Q-values (1 % FDR). Data were processed using R (version 4.1.1) and functions implemented in the pipeline of limma (version 1.30.7). The volcano plots were visualized using ggplot2 and the Gene Ontology analysis was performed using Metascape.

#### Targeted proteomics of proteotypic dermokine peptides

Peptides (500 ng and 0.25 pmol for each heavy peptide) from *DMKN* αβ^−/−^, *DMKN* βγ^−/−^ and WT 3D organotypic skin cultures were loaded onto a 2 cm C18 trap column (Thermo Fisher Scientific), connected in-line to a 50 cm C18 reverse-phase analytical column (Thermo Fisher Scientific) using 100% Buffer A (0.1% Formic acid in water) at 750 bar, using the EasyLC 1200 HPLC system (Thermo Fisher Scientific), and the column oven operating at 30 °C. Peptides were eluted over a 70-minute gradient ranging from 10% to 95% of 80% acetonitrile, 0.1% formic acid at 250 nL/min, and the Q-Exactive instrument (Thermo Fisher Scientific) was run in scheduled parallel reaction monitoring (PRM) mode for the *DMKN* βγ^−/−^ 3D organotypic skin cultures. Full MS spectra were collected at a resolution of 70000, with an AGC target of 3e^6^ or maximum injection time of 10 ms and a scan range of 350–1400 m/z. The MS^2^ spectra were obtained at a resolution of 35000, with an AGC target value of 1e^6^ or maximum injection time of 128 ms, a normalised collision energy of 27. For the *DMKN* αβ^−/−^ 3D organotypic skin cultures the MS^2^ spectra were obtained at a resolution of 17500. MS performance was verified for consistency by running complex cell lysate quality control standards, and chromatography was monitored to check for reproducibility.

Skyline 22.2.0.351 was used to analyze the PRM data. Target peptides were imported after setting filter parameters at MS^1^ precursor mass analyzer to orbitrap, resolving power of 70000 at 400 m/z and MS^2^ acquisition method set to PRM orbitrap resulting power of 30000 at 400 m/z. Endogenous peptides had a fixed modification of carbamido-methylation of cysteines (+57.021 Da). In addition, heavy peptides had fixed modifications of heavy lysine and heavy arginine amino acids on their C-terminus (+8.014 and +10.008 Da). Both an *in-silico* spectral library was generated via Prosit^64^ and a manual spectral library was created using the same instrument setup in DDA mode. The dermokine-proteotypic peptide – LGFINWDAINK – was removed from data analysis as well as - FGTNTQGAVAQPGYGS. The transition peak areas and their boundaries were manually adjusted and derived from the heavy peptide retention times. Endogenous peptides were normalized based on their ratio to heavy peptides. Final calculations were based on the sum of transition areas. The log_2_ transformed fold changes of summed total dermokine transition areas were calculated.

#### Mass spectrometry-based DIA phosphoproteomics and proteomics

*DMKN* αβ^−/−^, *DMKN* βγ^−/−^ and WT keratinocytes were washed three times with PBS (PAN Biotech) and incubated in DMEM/F-12 (Thermo Fisher Scientific) without FBS for 24 h. *DMKN* αβ^−/−^, *DMKN* βγ^−/−^ and WT keratinocytes were either incubated with buffer (25 mM Tris-HCl, 100 mM glycine, pH 7.3, 10% glycerol) or treated with recombinant dermokine (50 ng/mL) for 1 h. Cells were scraped from the dishes and the cell suspension was transferred into a falcon tube and washed three times before frozen in liquid frozen in liquid nitrogen. After thawing, cells were immediately resuspended in 8 M Urea and 50 mM Tris-HCl (pH 8.0). 200 µg lysate, determined by a BCA kit (Thermo Fisher Scientific), per sample were reduced with dithiothreitol (DTT) (Sigma-Aldrich) (1 mM final concentration), alkylated by iodoacetamide (Sigma-Aldrich) (5 mM final concentration), and digested with Lys-C (FUJIFILM) in an enzyme/protein ratio of 1:100 (w/w) for 2 h, shaking, followed by an 8-fold dilution with 50 mM Tris-HCl (Sigma-Aldrich) (pH 8.0) to 1 M Urea (Sigma-Aldrich) and further digested overnight with trypsin (Promega) 1:100 (w/w). Protease activity was quenched by acidification with TFA (Sigma-Aldrich) to a final concentration of 1% and peptides were purified by SPE using HR-X columns (Macherey-Nagel) in combination with C18 cartridges (Macherey-Nagel). The peptides were frozen in liquid nitrogen and lyophilized prior to phosphopeptide enrichment. The enrichment of phosphopeptides was automatically performed on a Bravo Automated Liquid Handling Platform (Agilent). The Fe (III)-NTA cartridges (Agilent) (5 µL) were initially primed with 200 µL 0.1% TFA (Sigma-Aldrich) in acetonitrile and then balanced with 0.1% TFA in 80% acetonitrile (Thermo Fisher Scientific) (equilibration/washing buffer). Peptides were redissolved in 200 µL of equilibration buffer and loaded onto the cartridges at a flow rate of 5 µL/min. Cartridges were then rinsed twice with 200 µL of washing buffer at a flow rate of 10 µL/min. Phosphopeptides were extracted first with 50 µL of 1% ammonia followed by 50 µL of 1% ammonia in 80% acetonitrile (Thermo Fisher Scientific) at a flow rate of 5 µL/min into 5 µL of 99% formic acid (Thermo Fisher Scientific). These samples were then concentrated via a lyophilizer and reconstituted in 20 µL of 0.1% formic acid (Thermo Fisher Scientific) for subsequent LC-MS/MS analysis.

Both the proteome and phosphoproteome of each condition, collected in triplicates, have been acquired using HRMS^1^-DIA methods with a mass range from either 400 – 1000 m/z and 58 min for the proteome or 400 – 1400 m/z for the phosphoproteome and 140 min.

For each resuspended proteome (200 ng), peptides were analysed using the ‘20 samples per day’ method on the EvoSep One instrument (EvoSep). Peptides were eluted over a defined 58-min gradient and analysed by an Orbitrap Exploris 480 mass spectrometer (Thermo Fisher Scientific), with following settings: Spray voltage was set to 2.3 kV, funnel RF level at 40, and heated capillary at 240 °C. Full MS spectra were collected at a resolution of 120000, with an AGC target of 300% or maximum injection time set to ‘auto’ and a scan range of 400–1000 m/z. The MS^2^ spectra were obtained in DIA mode in the Orbitrap operating at a resolution of 30000, with an AGC target 1000% or maximum injection time set to ‘auto’, a normalised HCD collision energy of 32. The isolation window was set to 8 m/z with a 1 m/z overlap and window placement on. Each DIA experiment covered a range of 200 m/z resulting in three DIA experiments (400-600 m/z, 600-800 m/z and 800-1000 m/z). Between the DIA experiments a full MS scan was performed. MS performance was verified for consistency by running complex cell lysate quality control standards, and chromatography was monitored to check for reproducibility.

Each resuspended enriched keratinocyte phosphoproteome (1 µg) was analysed with an Easy-nLC 1200 (Thermo Fisher Scientific) coupled to an Orbitrap Exploris 480 mass spectrometer (Thermo Fisher Scientific) operated with a DIA method; 5 µL of each sample was injected. Peptides were separated on a fused silica HPLC-column tip (I.D. 75 µm, self-packed with ReproSil-Pur 120 C18-AQ, 1.9 µm (Dr. Maisch) to a length of 20 cm) using a gradient of mobile phase A (0.1% formic acid in water) and mobile phase B (0.1% formic acid in 80% acetonitrile in water). The column temperature was maintained at 50 °C using the EASYSpray oven. The total gradient time was 140 min and went from 6% to 23% acetonitrile (ACN) in 95 min, followed by 30 min to 38%. This was followed by 5 min increase to 60% ACN, and a washout by 10 min increase to 90% ACN. Flow rate was kept at 250 nL/min. Re-equilibration was done prior to sample pickup and prior to loading with a volume of 4 µL of 0.1% FA buffer at a maximum pressure of 750 bar. Spray voltage was set to 1.9 kV, funnel RF level at 40, and heated capillary at 275 °C. Full MS spectra were collected at a resolution of 120000, with an AGC target of 300% or maximum injection time set to ‘auto’ and a scan range of 400–1400 m/z. The MS^2^ spectra were obtained in DIA mode in the Orbitrap operating at a resolution of 60000, with an AGC target 1000% or maximum injection time set to ‘auto’ and a normalised HCD collision energy of 32. The isolation window was set to 6 m/z with a 1 m/z overlap and window placement on, resulting in 175 DIA windows. Each DIA experiment covered a range of 200 m/z resulting in three DIA experiments (400-600 m/z, 600-800 m/z, 800-1000 m/z, 1000-1200 m/z and 1200-1400 m/z). Between the DIA experiments, a full MS scan was performed. All data were acquired in profile mode using positive polarity, application mode was set to peptide, FAIMS mode was not installed, advanced peak determination was set to true, and default charge state was set to 2. MS performance was verified for consistency by running complex cell lysate quality control standards, and chromatography was monitored to check for reproducibility. The median peak width was 0.91 min, and the full width at half maximum was 0.53 min. Each MS^1^ window had 175 MS^2^ DIA windows, achieving a 5.17 s cycle time and 10 median data points per peak on MS^1^ level with a 140 min gradient length.

Two independent searches for the proteome and phosphopeptide samples were performed. For the proteome, raw files were analysed using Spectronaut™ (Biognosys, version 18.5) directDIA+ (deep) default settings, spectra were matched against the homo sapiens reference FASTA database (UP000005640; downloaded from Uniprot on 27th of December 2022). Dynamic modifications were set as Oxidation (M) and Acetyl on protein N-termini. Cysteine carbamidomethyl was set as a static modification. Protein quantitation was done on the MS^1^ level, data filtering set to Q-value and the data was normalized by RT-dependent local regression model and protein groups were inferred by IDPicker. For phosphopeptides, raw files were analysed using Spectronaut™ (Biognosys, version 17.4) directDIA+ (deep), spectra were matched against the homo sapiens database (UP000005640; downloaded from Uniprot on 27th of December 2022) with the BGS Phospho PTM workflow with the following modifications: deamidation (NQ) and oxidation (M) were added as dynamic modifications together with Phospho (STY) and Acetyl on protein N-termini. Cysteine carbamidomethyl was set as a static modification. The proteomics dataset was run-wise imputed via Spectronaut™ (Biognosys). All results were filtered to a 1% FDR, data filtering set to Q-value and protein quantitation done on the MS^1^ level. The data was cross-normalized by a RT-dependent local regression model on modified (phospho)-peptides. The PTM probability cutoff was set to 0.

The phosphoproteomic data analysis was performed using modified function from the “PhosR” package embedded in custom scripts in R (version 4.3.2). Data at protein level was processed using R (version 4.3.2). The Supplemented WT samples were not considered for any further analysis due to the high coefficient of variation. Due to the nature of DIA, all peptides were acquired, recording single low-intense phosphorylated precursors (singletons)^65^. To remove singletons, we excluded phosphorylated peptides that contained low quality chromatography peaks (>0.6), utilizing Spectronaut’s ability to evaluate the shape of chromatographic peaks (FG.ShapeQualityScore (MS^1^)). We then imported the filtered phosphorylated peptides into MaxQuant and assigned site localization probabilities.

For the analysis only peptides with a site localization probability threshold of at least 0.75 were considered in the bioinformatic analysis^31^. Next, we excluded phosphorylated sites that were less than 70% present in at least one condition. Finally, we removed phosphorylated sites that contained any missing value after removing all phosphorylated sites with any missing values we ended up with 4453 fully quantifiable phosphorylated sites, including 299 tyrosine phosphorylated sites without specific enrichment methods. For kinase-substrate enrichment analysis using data shown in Figure 3, the filtered Spectronaut™ (Biognosys, version 18.5) reports were transformed into modification-specific peptide-like reports using the plugin peptide collapse in Perseus (version 2.0.9.0) with “EG.PTMAssayProbability” as grouping column. A localization cutoff of 0.75 was applied and the same variable PTMs as listed above, and summed intensities were log_2_-transformed. No imputation was performed in the phosphoproteomics dataset. The limma package was employed to calculate the fold change as well as the moderated p-value for the comparisons between genotypes for both the phosphorylated peptides and the log_2_-transformed protein abundance values. To analyze the proteome, we used the limma package in R and the volcano plots were plotted using ggplot2. Gene ontology analysis was performed using Metascape. The phosphoproteomics datasets were aligned with upstream kinases by inferring kinase activity through clustering phosphorylated sites based on either their known kinase targets (profile score) or their common phosphorylation motifs (motif score), following the Kinase-Substrate prediction approach^32,66^. In brief, phosphorylated sites were z-transformed and mapped to known kinases using PhosphoSitePlus® data^32^. Profile and motif scores were calculated, averaged, and used to determine a combined score for each site^32^. The combined scores were visualized in a heatmap and was clustered as circular signalome map^32^. The kinase activity score is computing the kinase-substrate profile for each kinase by calculating the median phosphosite quantification for kinase in the quantification matrix^32^. The kinases and their respective kinase activity score were tested for significance (one-way ANOVA) among all genotypes. Clusters from the circular signalome map were subjected to GO analysis, including all ontologies and showing the top 3 GO terms, via the clusterProfiler package^32^.

To normalize the intensity of the phosphorylated peptide we divided the raw intensities of the phosphorylated sites through the respective protein quantities^39^. All Figures were plotted in R using ggplot2.

#### Cell lysis and SDS-PAGE immunoblotting

Cells from endogenous *DMKN* βγ^−/−^, *DMKN* αβ^−/−^ and WT keratinocytes (Figure 4c) as well as supplemented *DMKN* βγ^−/−^, *DMKN* αβ^−/−^ and WT keratinocytes (Figure S9a) were washed in ice-cold PBS once before being scraped into lysis buffer (25 mM HEPES pH 7.4, 150 mM KCl, 2 mM MgCl2, 1 mM EGTA, 0.5% (v/v) Triton X-100) supplemented with EDTA-free Protease Inhibitor Cocktail (Sigma-Aldrich), PhosSTOP (Roche), and 1 µM DTT. Lysates were cleared by centrifugation at 13000 rpm at 4 °C for 10 minutes. Supernatant was transferred to fresh tubes, concentrations adjusted using Bradford assay (BioRad), and proteins were denatured by adding Laemmli sample buffer (LSB) followed by heating for 5 minutes at 70 °C. Samples were loaded on 9 % Bis-Tris gels and run at 150 V in Tris running buffer or on 4-12 % Bis-Tris gels (Invitrogen) and run in MOPS buffer (Invitrogen). Proteins were transferred onto 0.2 µM nitrocellulose membranes (BioRad) using the Trans-Blot Turbo System (BioRad), and membranes were blocked for 30 min in 5 % milk in TBS-T. After blocking, membranes were incubated with primary antibodies in 3% BSA-TBS-T overnight at 4 °C while shaking. After washing in TBS-T for 5 minutes, membranes were incubated with HRP-coupled secondary antibodies for 45 minutes at room temperature. After 3 x 5 minutes TBS-T washes, protein bands were developed using ECL Western Blotting Substrate (BioRad). Chemiluminescence was detected using BioRad’s ChemiDoc Imaging Systems and software.

#### Transfection and RNA interference

Cells were cultured in DMEM/F12 (Thermo Fisher Scientific) with Lipofectamine (Invitrogen) according to the manufacturer’s instructions, and all assays were conducted 72 h after transfection. Double-stranded, validated Stealth siRNA oligonucleotides targeting human p120 human (HSS175607 (5’-CAGCUCGAGGCUAUGAGCUCUUAUU-3’), HSS102463 (5’-GGCUAGAGGAUGACCAGCGUAGUAU-3’) and HSS102464 (5’-GCAGCUCCCAAUGUUGCCAACAAUA-3’), called siRNA#1, siRNA#2 and siRNA#3 respectively) were purchased from Invitrogen. As a negative control, duplexes with irrelevant sequences (5’-GGGAUACCUAGACGUUCUA-3’, called scrambled siRNA) were purchased from Sigma-Aldrich. *DMKN* βγ^−/−^, *DMKN* αβ^−/−^ or WT keratinocytes were transfected either with siRNA#1, siRNA#2, siRNA#3, scrambled siRNA or lipofectamine alone. Immunoblotting, as described previously, of *DMKN* βγ^−/−^, *DMKN* αβ^−/−^ or WT keratinocytes transfected either with siRNA#1, siRNA#2, siRNA#3 or lipofectamine alone (control) was performed to test for p120 expression.

#### Cell-cell adhesion assay

This assay was performed as described before^40^. Briefly, cells from *DMKN* βγ^−/−^, *DMKN* αβ^−/−^ and WT keratinocytes were detached using Trypsin-EDTA (Thermo Fisher Scientific), keeping both calcium dependent and independent cell adhesion molecules intact. Cells were washed with PBS counted and resuspended in DMEM/F12 (Thermo Fisher Scientific) containing 25 mM HEPES and 1 mM CaCl_2_ to a concentration of 2e^5^ cells/mL. 10 mL of each cell suspension was placed in the respective non-adhesive 100 mm dish (VWR), as previously described^40^. The dishes were placed in an agitating incubator (Infors AG) at 37 °C while shaking (30 rpm) for 100 minutes. Immediately after 25 images with an EVOS M5000 (Thermo Fisher Scientific) per dish were automatically captured in a counterclockwise manner. Clusters of more than 4 cells from each image from each replicate were counted^40^ and analyzed via GraphPad Prism (Dotmatics). A one-way ANOVA was used to detect changes among the conditions and illustrated as violin plots.

As described previously, *DMKN* βγ^−/−^, *DMKN* αβ^−/−^ or WT keratinocytes were transfected either with siRNA#1, siRNA#2, siRNA#3 or scrambled siRNA in 6 well plates for 72 hours. The transfected cells were dissociated using Trypsin-EDTA (Thermo Fisher Scientific) and treated as described previously. Immediately after the assay, 4 randomly chosen images were captured per dish to capture clusters of more than 4 cells. Cell cluster (≥4) were counted, summarized per replicate and each siRNA transfected *DMKN* αβ^−/−^, *DMKN* βγ^−/−^, WT keratinocyte replicate was normalized to the average WT cell cluster number. Subsequently all normalized siRNA cell cluster values were averaged for each genotype. A two-way ANOVA and a Fishers Least Significant Difference test was used to show changes among the conditions.

#### Immunofluorescence and quantification of patient tissue

Formalin-fixed paraffin-embedded (FFPE) tissue samples from SCCs, wounds and psoriasis were selected and retrieved from the SKINTEGRITY.CH biobank of the Department of Dermatology, University Hospital Zurich. FFPE blocks of healthy skin samples were also included. The blocks were used to prepare 7 µm tissue sections to be subsequently stained. Patient information is available in original data file.

FFPE patient tissue sections were dried for 30-60 min at 60 °C before being dewaxed, rehydrated, and buffered in PBS. Antigen retrieval was performed with 0.1 M sodium citrate buffer (pH 6) for 45 minutes at 95 °C. After three washes with PBST (PBS + 0.1% Tween), unspecific binding to tissue sections was blocked for 1 hour at room temperature with 12% BSA in PBST. Primary antibody was diluted (catenin δ-1 (Cell Signalling Technology) (1:200), dermokine (Abcam) (1:250), phospho-catenin δ-1 Tyr228 (Abcam) (1:50), phospho-catenin δ-1 Thr310 (Abcam) (1:250) and phospho-catenin δ-1 Ser252 (Cell Signalling Technology) (1:250)) in blocking buffer and incubated overnight at 4 °C. After three washes with PBS, slides were incubated for 1 h at room temperature with the secondary antibody (anti-rabbit-IgG-Cy3, 1:200) and 1 µg/ml Hoechst 33342 (Sigma-Aldrich) in blocking buffer. After another three washes with PBS, sections were mounted using Mowiol with DABCO®. Slides were scanned with an Axioscan 7 slide scanner (Carl Zeiss AG).

Staining intensity was quantified using Qupath (version 0.5.0). Positively stained area was determined using a pixel thresholder and median intensity of stained area was reported. Data was analyzed in GraphPad Prism (Dotmatics). A one-way ANOVA with post hoc Fishers LSD test or a one-tailed, unpaired *t* test (healthy skin compared to non-healing wounds) was applied to detect changes among conditions.

#### Quantification and statistics analysis

All experiments were repeated at least three times as independent biological replicates with similar results. Representative Western blots or IF or IHC images are shown. See figure legends for statistical details. The number of independent experiments, treatments and relative controls and statistical analysis are indicated in Figure legends.

#### Ethics declaration

All patient-derived samples used were surplus material from routine surgeries and were provided by the Dermatology Department of the University Hospital Zürich, Switzerland, with the assistance of the SINTEGRITY.CH biobank. The surplus biopsies stored in the biobank were from consenting patients. The use of material for research purposes was approved by the local ethics commissions (Cantonal Ethic Commission Zürich: project no. 2017-00684). All experiments conformed to the principles set out in the World Medical Association’s Declaration of Helsinki and the Department of Health and Human Services Belmont Report.

## Acknowledgement

The authors would like to thank Prof. Dr. Sabine Werner for her continuous support, especially after U.a.d.K.’s sudden passing. We apologize to authors whose work is not cited owing to space constraints. C.F. has received funding from the Wellcome Trust (107636/ Z/15/Z and 107636/Z/15/A) and the Novo Nordisk Foundation Young Investigator Award (call 2022, NNF22OC0070845). U.a.d.K. was supported by a Novo Nordisk Foundation Young Investigator Award (call 2016, NNF16OC0020670) and received funding from the LEO Foundation (LF-OC-19-000033 and LF-OC-23-001220). V.C. is supported by the LEO Foundation (LF-OC-23-001220). We would like to thank Skintegrity.CH. Mass spectrometry analysis was performed at the Proteomics Core, Technical University of Denmark, and the Metabolomics and Proteomics Platform at the University of Fribourg. Human n/TERT keratinocytes were a kind gift from Prof. Dr. Edel O’Toole.

## Author Contributions

U.a.d.K. conceived the project and acquired funding. U.a.d.K., C.F., J.D. and V.C. designed the experiments. V.C. and W.T. performed the generation of *DMKN* βγ^−/−^ keratinocytes. V.C., T.A.B., A.M.H and S.M. generated the *DMKN* αβ^−/−^ keratinocytes. V.C. generated 3D organotypic skin cultures, all the omics experiments and performed data analysis. M.S. provided experimental guidance for semi-automated phosphoproteomics. Patient tissue biopsies were cut by G.R. T.W. stained tissue slides and microscopically captured the images. Images were analysed by V.C. and T.W. C.R.E. carried out phosphorylated p120 immunoblotting. V.C. performed RNA interference and cell-cell adhesion assays. C.C. provided experimental input for the knockdown experiments. C.F. and V.C. wrote the manuscript. All authors read, revised and approved the manuscript.

