## Supplemental Figures for "Multi-omics analysis of keratinocytes reveals dermokine-dependent regulation of cell-cell adhesion via p120"

**Supplementary Figure 1**

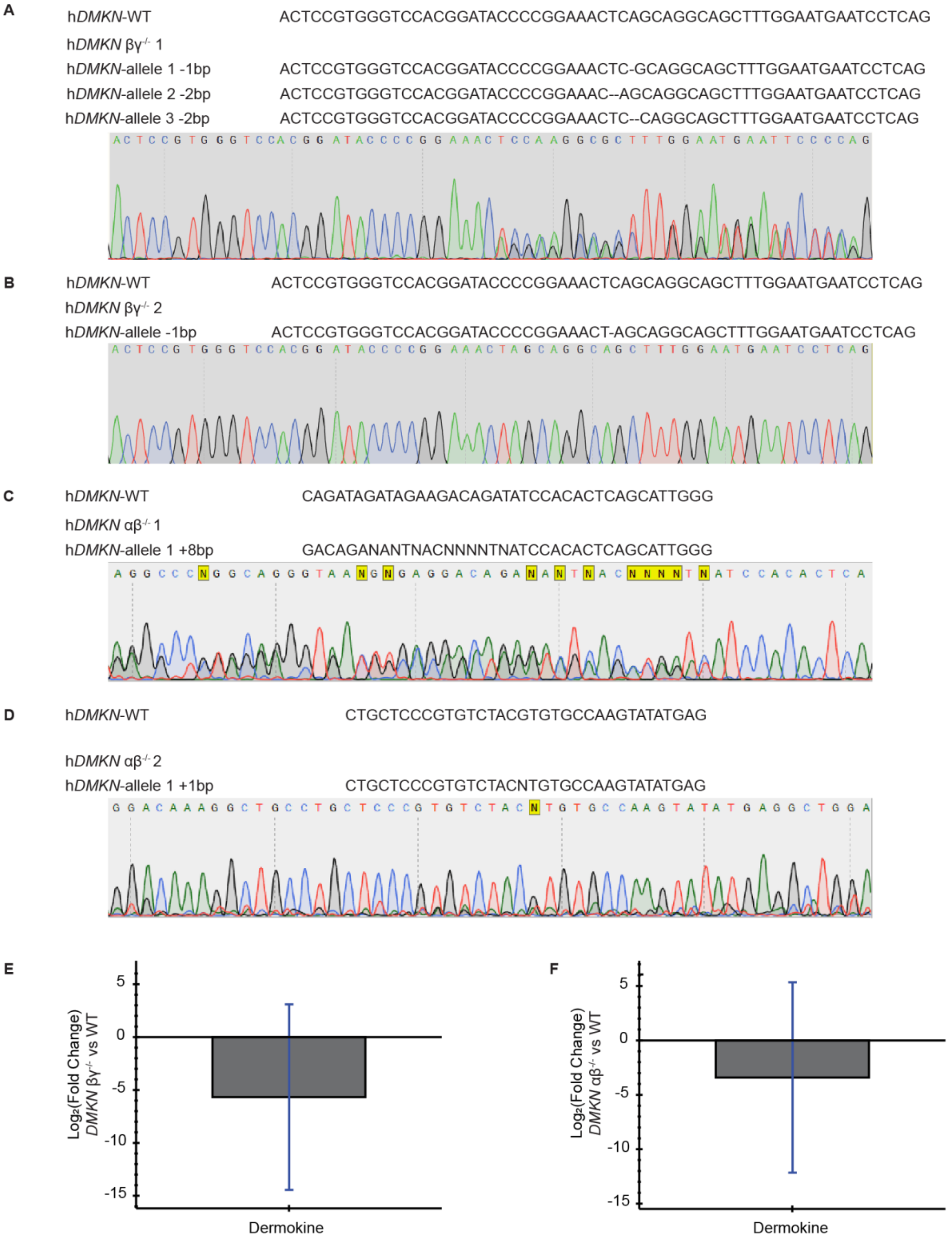


**Supplementary Figure 1. Sanger sequencing validates the ablation of human *DMKN* αβ or *DMKN* βγ from keratinocytes. (A and B)** Results of the sanger sequencing of the CRISPR/*Cas9* target sequence and the sanger sequence of *DMKN* $\beta\gamma^{-/-}$ clone 1 (**A**) or *DMKN* $\beta\gamma^{-/-}$ clone 2 (**B**) keratinocytes. **(C)** Sanger sequences of Chromosome 19 (GRCh 38.p14) region 35500725 to 35500763 are shown for both WT and *DMKN* $\alpha\beta^{-/-}$ keratinocytes clone 1 illustrating indel (insertion/deletion) sites. **(D)** The sanger sequence of region 35501517 to 35501551 of Chromosome 19 (GRCh 38.p14) are shown for both WT and *DMKN* $\alpha\beta^{-/-}$ keratinocytes clone 12. Both (**C**) and (**D**) show the presence of indels after the excision of exon 17. **(E and F)** Targeted proteomics showing the quantitative measurement of proteotypic dermokine peptides, in combination with isotopically labelled proteotypic peptides, in *DMKN* $\beta\gamma^{-/-}$ **(E)** or *DMKN* $\alpha\beta^{-/-}$ **(F)** compared to WT 3D organotypic skin cultures. Endogenous peptides were normalized based on heavy isotopically labelled peptides. Values are the $Log_2$ transformed fold changes and negative $Log_{10}$ transformed false discovery rate adj. p values of three different 3D organotypic skin cultures.

**Supplementary Figure 2**

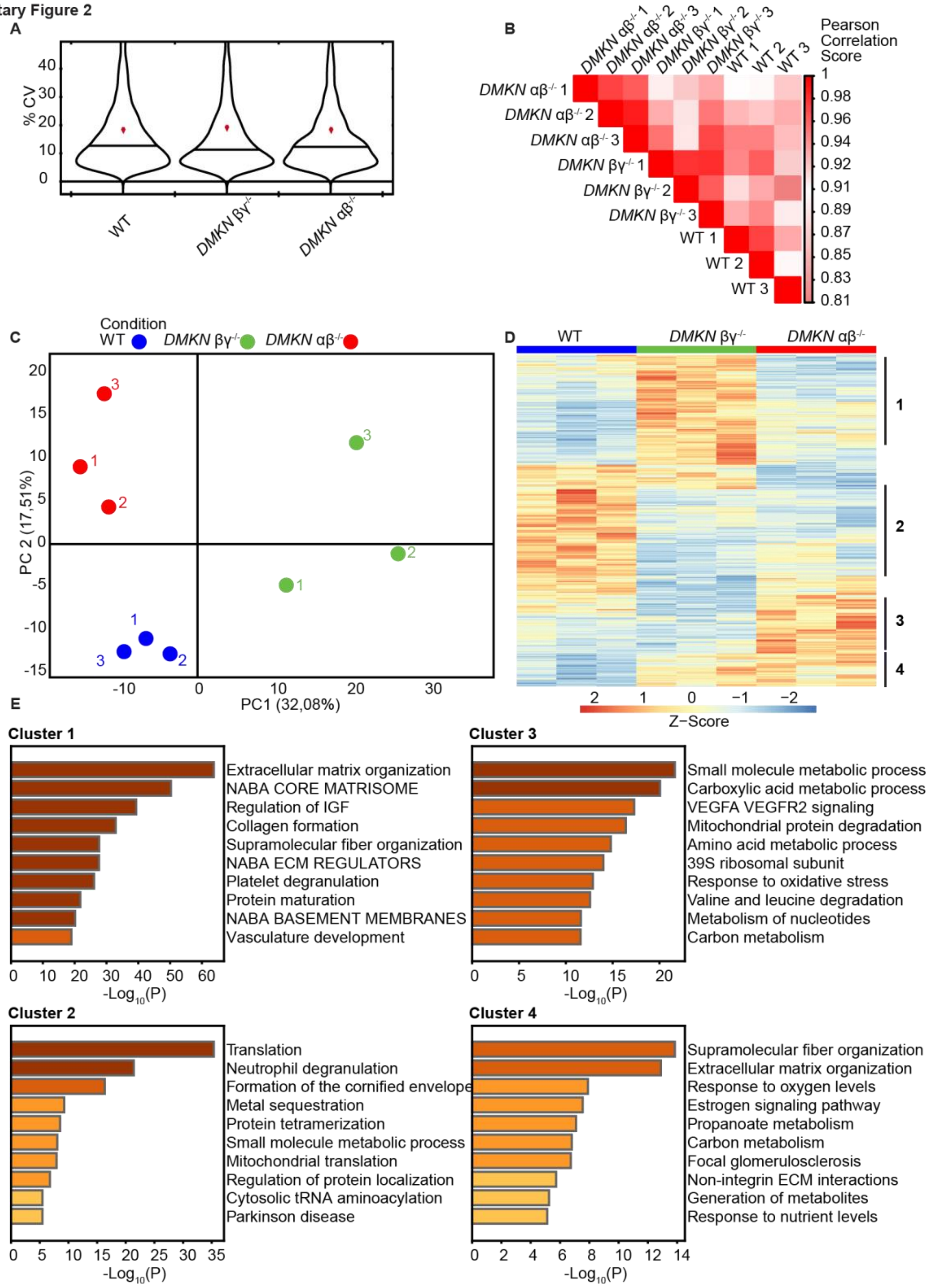


**Supplementary Figure 2: Proteomics analysis of the WT, *DMKN* αβ$^{-/-}$ and *DMKN* βγ$^{-/-}$ 3D organotypic skin cultures demonstrates high data quality. (A)** Coefficient of variation (CV) of the WT, *DMKN* αβ$^{-/-}$ and *DMKN* βγ$^{-/-}$ 3D organotypic skin culture proteomes. **(B)** Pearson **c**orrelation analysis of WT, *DMKN* αβ$^{-/-}$ or *DMKN* βγ$^{-/-}$ 3D organotypic skin cultures proteomes. **(C)** Principal component analysis (PCA) of the indicated data sets. **(D)** Differences in the proteomes of *DMKN* αβ$^{-/-}$, *DMKN* βγ$^{-/-}$ or WT 3D organotypic skin cultures visualized as heatmap and four clusters numbered 1 to 4 on the right. **(E)** Gene Ontology enrichment analysis of the different *DMKN* KO proteome clusters shown in **(D)**.

**Supplementary Figure 3**

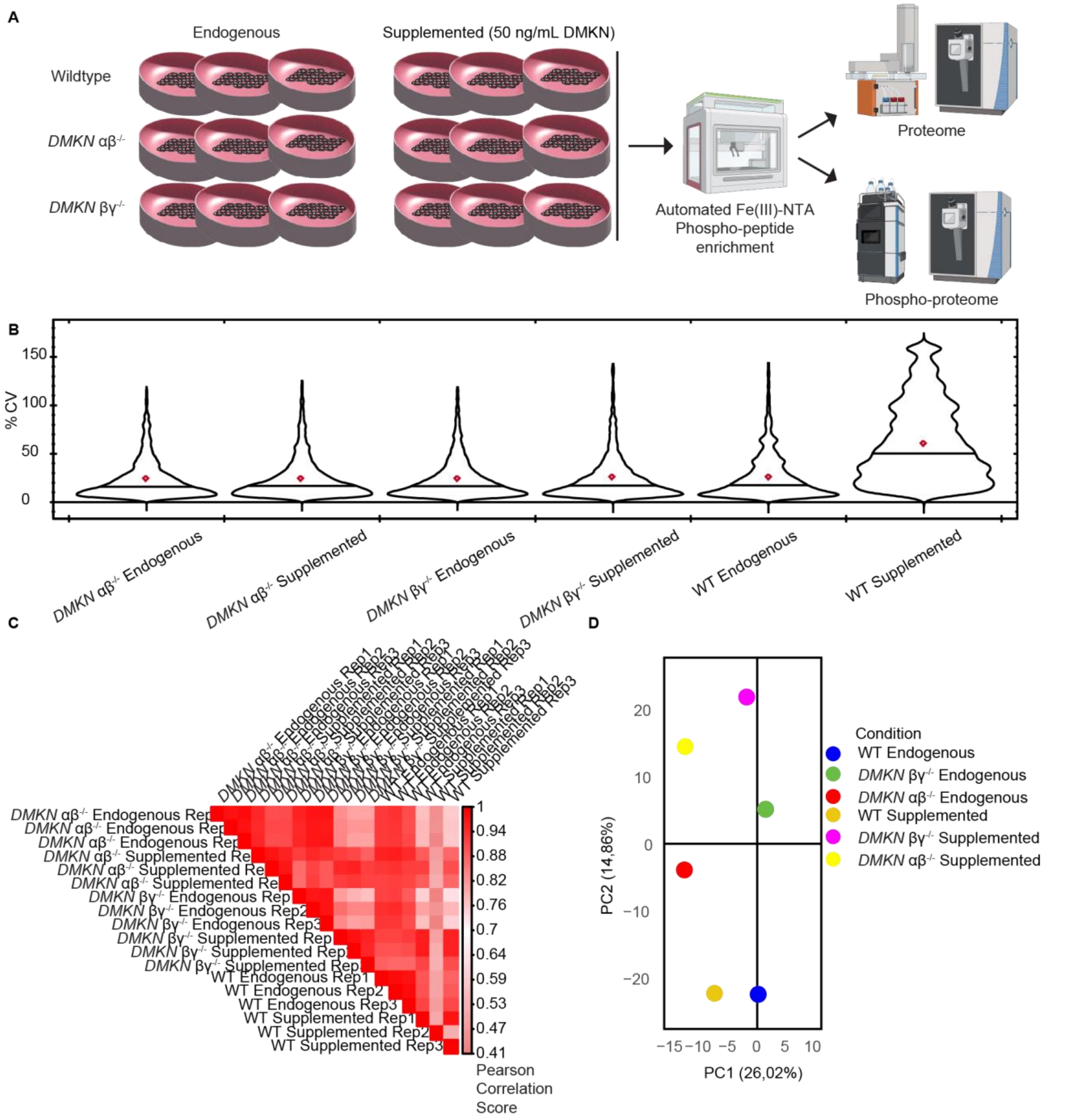


**Supplementary Figure 3: Evaluation of proteomics analysis of WT, *DMKN* αβ$^{-/-}$ and *DMKN* βγ$^{-/-}$ keratinocytes dataset detects changes among conditions (A)** Overview of the experimental setup which includes endogenous *DMKN* αβ$^{-/-}$ or *DMKN* βγ$^{-/-}$ and WT keratinocytes together with supplemented *DMKN* αβ$^{-/-}$ or *DMKN* βγ$^{-/-}$ and WT keratinocytes. The data was acquired on an Exploris Orbitrap with a 140-minute gradient in an optimized HRMS$^1$-DIA mode. **(B)** Coefficient of variation (CV) of endogenous and supplemented *DMKN* αβ$^{-/-}$ or *DMKN* βγ$^{-/-}$ and WT keratinocytes. **(C)** Correlation analysis of either endogenous or supplemented WT, *DMKN* αβ$^{-/-}$ or *DMKN* βγ$^{-/-}$ proteomes. **(D)** PCA of the median of each replicate within the indicated data sets.

Supplementary Figure 4

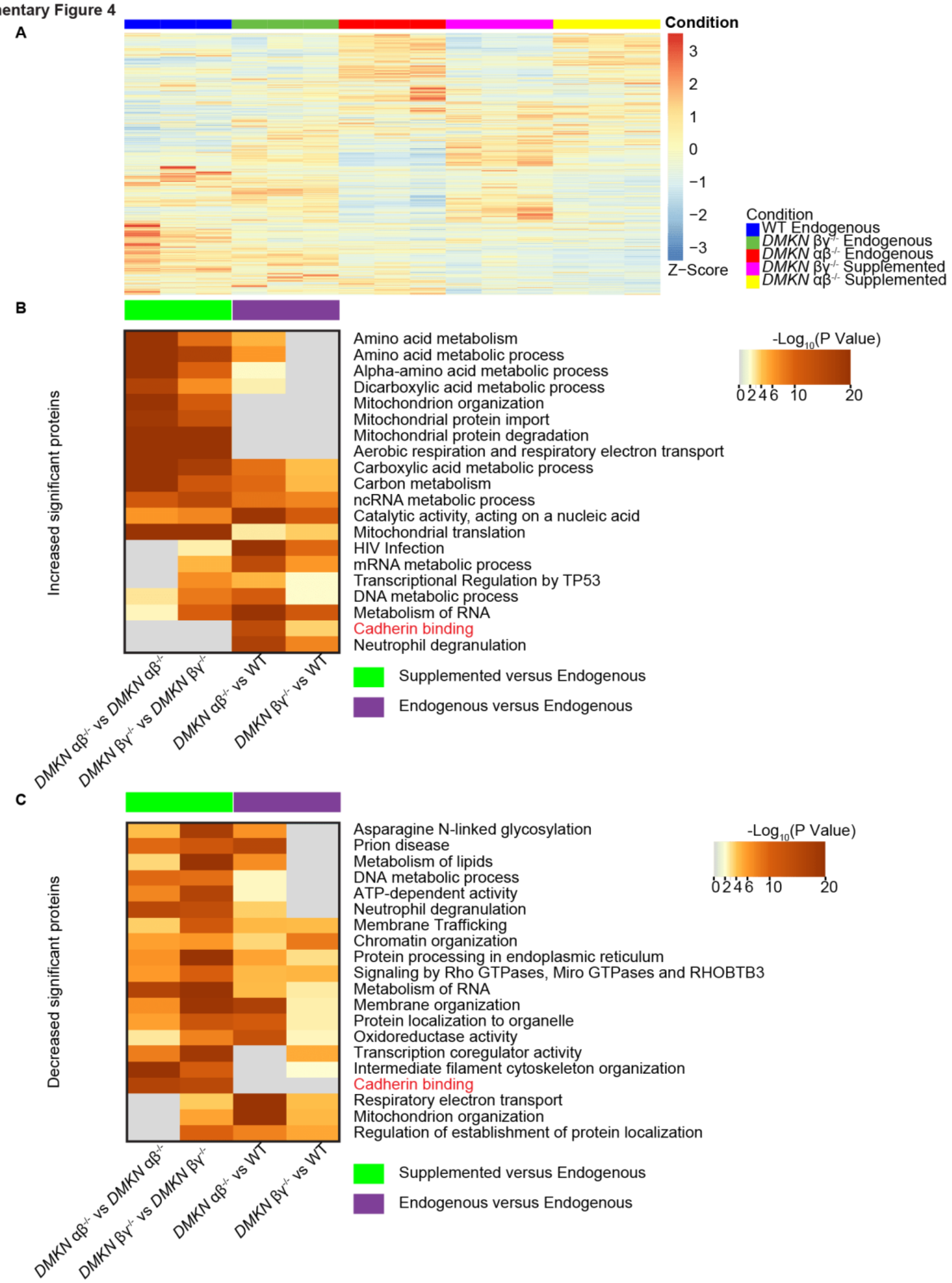


**Supplementary Figure 4: Pathway analysis of WT, *DMKN* αβ$^{-/-}$ and *DMKN* βγ$^{-/-}$ proteomes reveals proteins linked to cadherin binding in dermokine-ablated keratinocytes. (A)** Scaled proteome analysis of either supplemented or endogenous *DMKN* αβ$^{-/-}$ and *DMKN* βγ$^{-/-}$ or endogenous WT keratinocyte proteomes. **(B and C)** Gene ontology analysis of the *DMKN* βγ$^{-/-}$, *DMKN* αβ$^{-/-}$ and WT keratinocyte proteome reveals which pathways were significantly increased (**B**) or decreased (**C**) in supplemented relative to endogenous *DMKN* βγ$^{-/-}$ or *DMKN* αβ$^{-/-}$.

Supplementary Figure 5

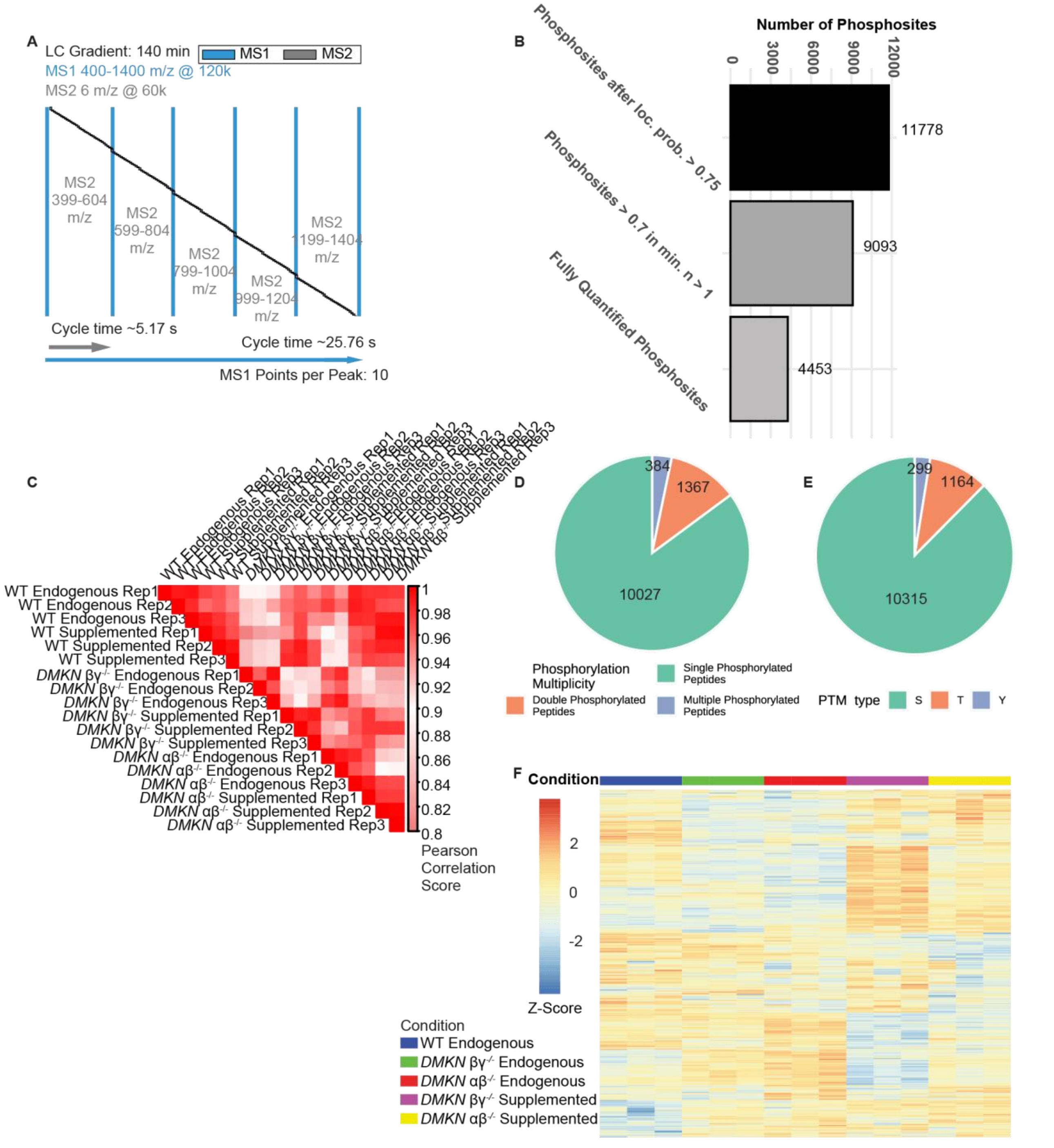


**Supplementary Figure 5: Quality assessment of the WT, *DMKN* αβ$^{-/-}$ and *DMKN* βγ$^{-/-}$ keratinocyte phosphoproteome dataset reveals changing phosphorylated proteins between WT and KO samples.** **(A)** Workflow of the method used to acquire the phosphoproteome. **(B)** Pearson correlation analysis of WT, *DMKN* αβ$^{-/-}$ or *DMKN* βγ$^{-/-}$ proteomes. **(C)** Distribution of singly doubly or multiple phosphorylated peptides. **(D)** Distribution of phosphorylated serine, threonine and tyrosine sites. **(B)** Total number of acquired phosphorylated sites after applying the site localization filter of 0.75 and excluding non-fully quantified phosphorylated sites. **(F)** Scaled phosphoproteome intensities showing differences between conditions.

Supplementary Figure 6

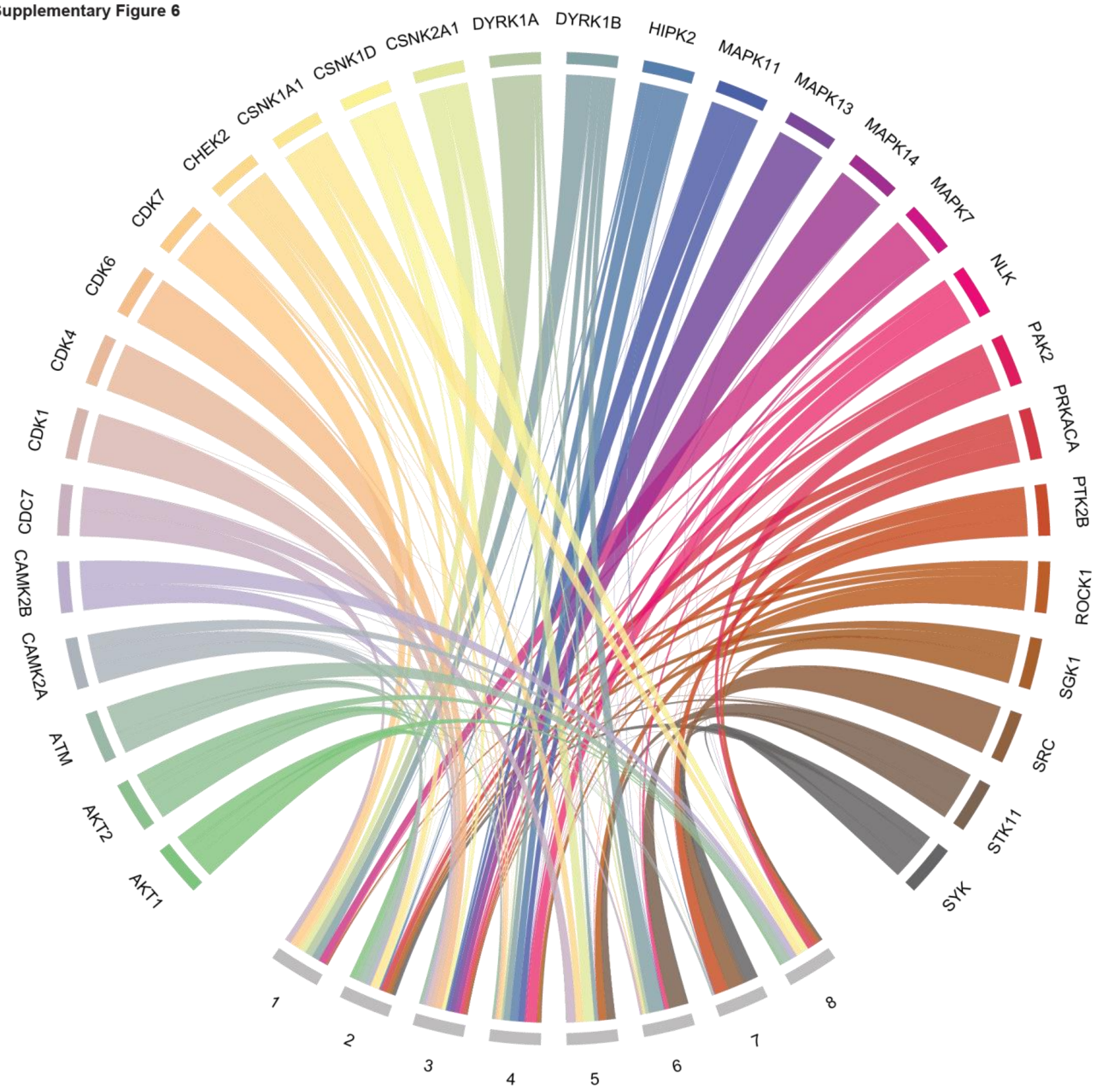


**Supplementary Figure 6: Clustering of phosphorylated proteins based on kinase preference shows that phosphorylated site modules share similar phosphorylation profiles.** Circular signalome map clustering phosphorylated sites according to kinase preference The calculated kinase-substrate scoring of phosphorylated sites demonstrate that sites cluster into phosphorylation modules. Cluster 5 and 7 contain phosphorylated tyrosine and threonine sites. Module 7 includes PTK2B, SRC and SYK activity.

Supplementary Figure 7

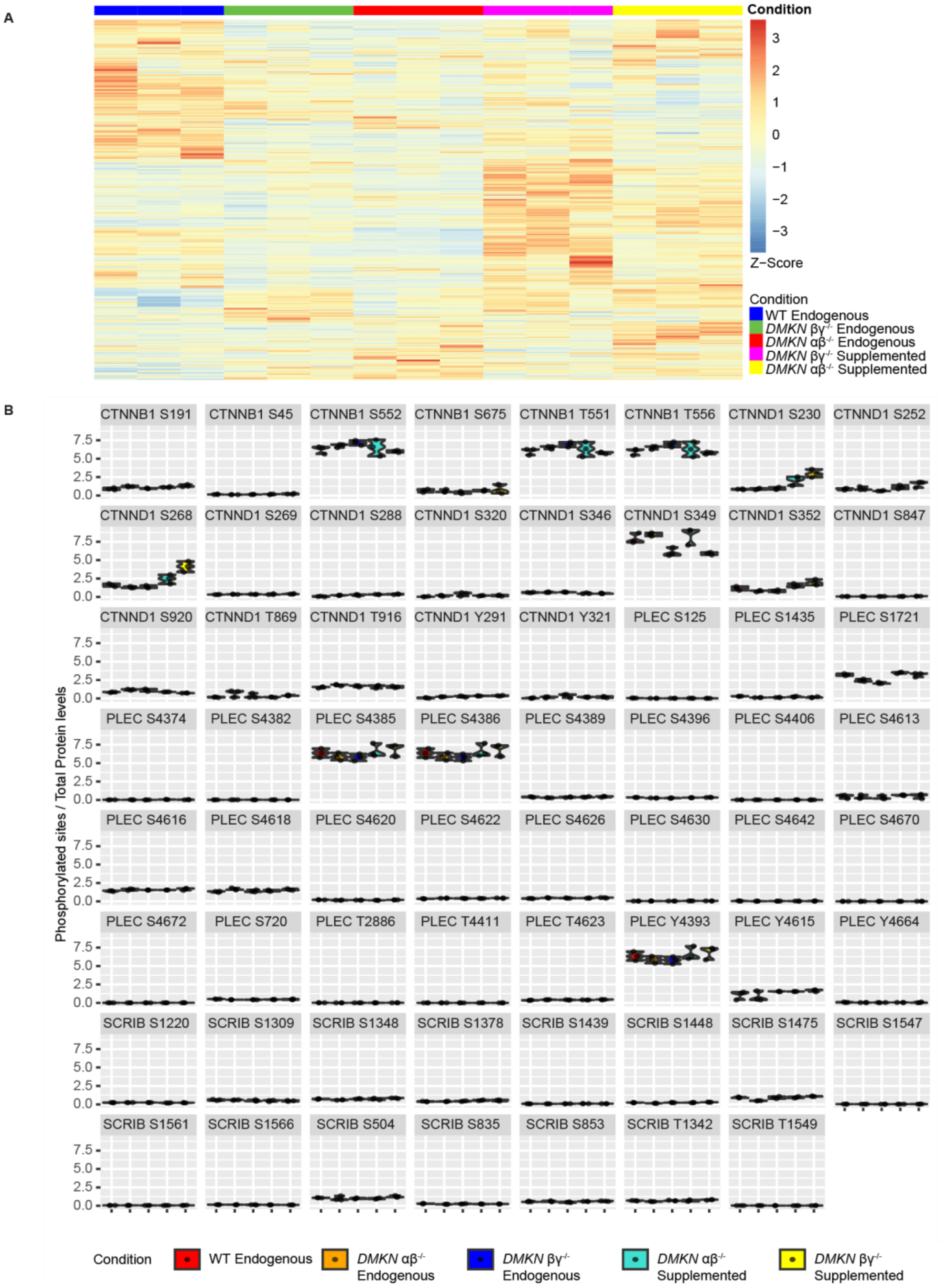


**Supplementary Figure 7: Normalized WT, *DMKN* αβ-/-, and *DMKN* βγ-/- keratinocyte phosphoproteome shows changes similar to not normalized phosphoproteome between the WT and dermokine KO samples. (A)** All phosphorylated proteins were normalized to their protein expression and scaled, demonstrating differences between conditions. **(B)** Selected phosphorylated sites normalized to protein levels.

Supplementary Figure 8

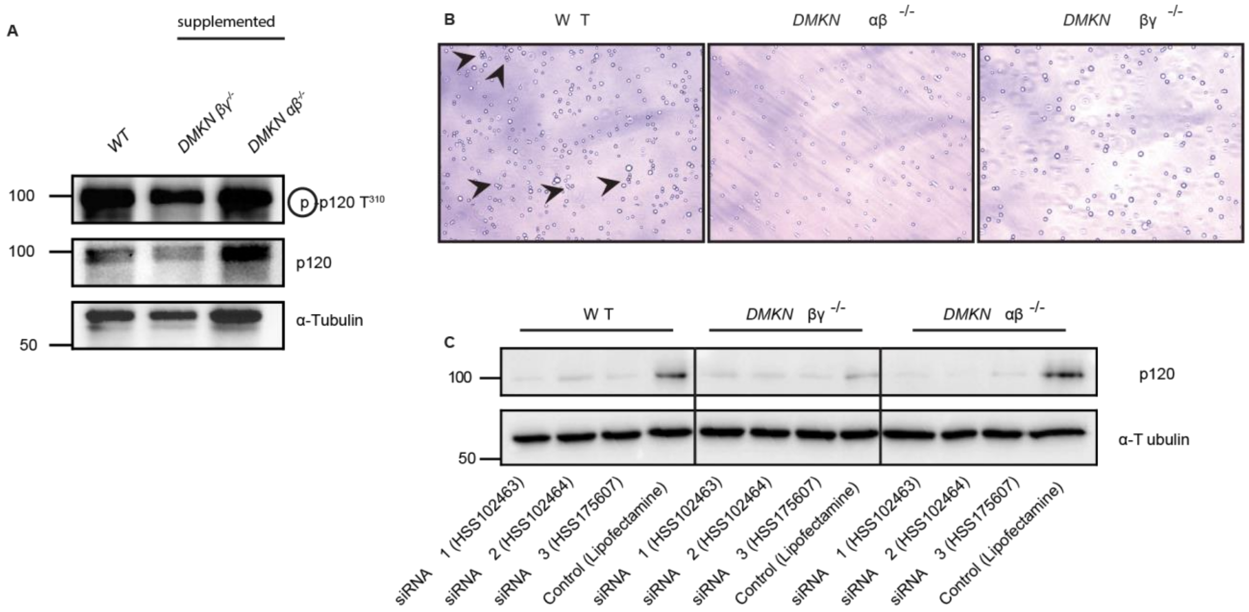


**Supplementary Figure 8: Validation of supplemented WT, *DMKN* αβ$^{-/-}$, and *DMKN* βγ$^{-/-}$ keratinocyte and functional assays shows dermokine**-mediated regulation of **cell-cell adhesion. (A)** Immunoblotting with indicated antibodies of WT and supplemented *DMKN* αβ$^{-/-}$, *DMKN* βγ$^{-/-}$ keratinocytes. **(B)** Representative images of the cell clusters from the cell-cell adhesion experiment quantified in Fig. 4D. Arrows indicate the presence of four or more cell clustering together. **(C)** Immunoblotting with the indicated antibodies of lysates from *DMKN* αβ$^{-/-}$, *DMKN* βγ$^{-/-}$ and WT keratinocytes incubated for 72 hours with p120 siRNA 1, 2 and 3 and a control (lipofectamine).

**Supplementary Figure 9**

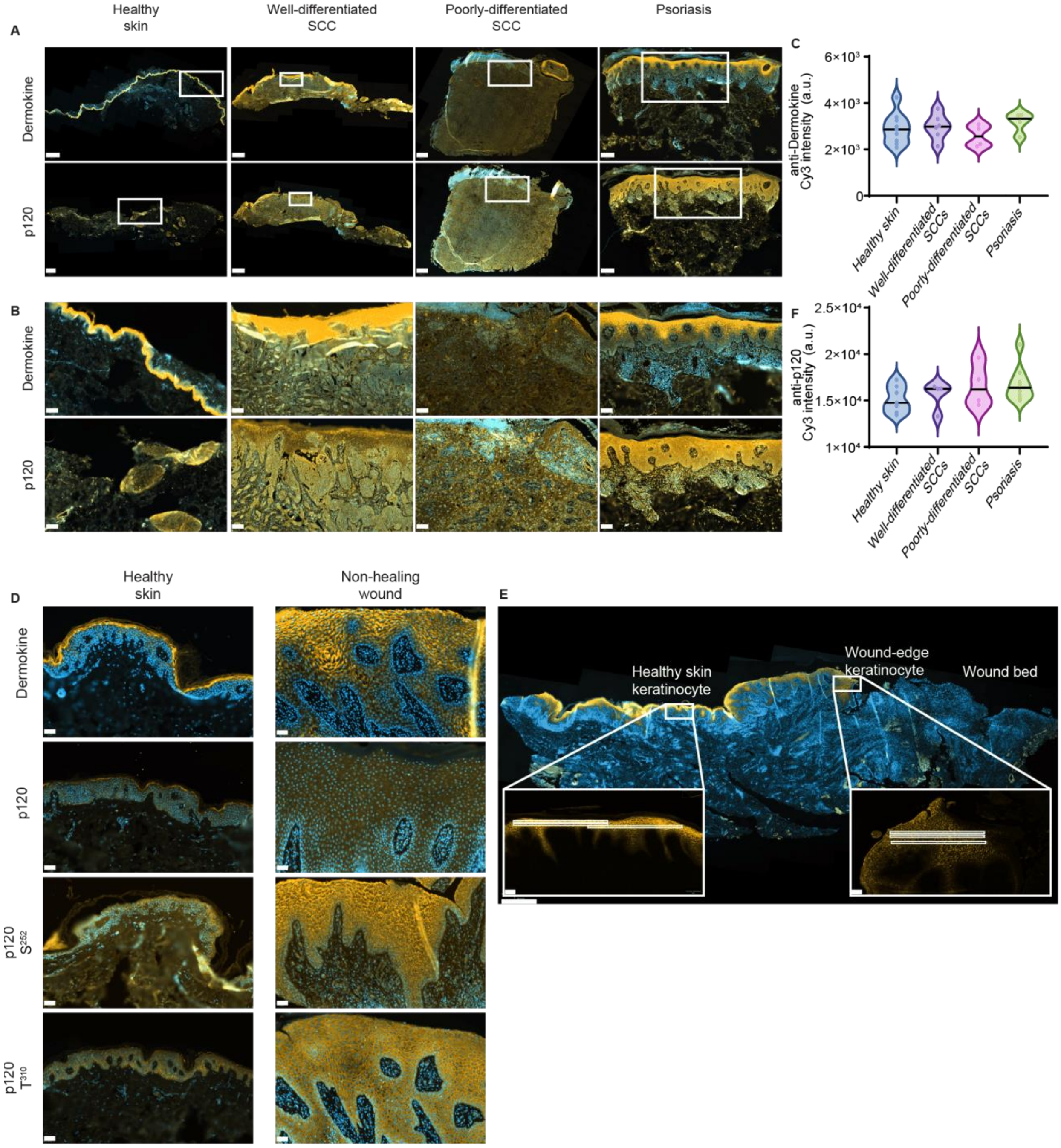


**Supplementary Figure 9: Immunofluorescence staining of healthy and psoriatic skin as well as poorly-differentiated squamous cell carcinoma (SCC) samples shows no change in p120 expression and wound-edge keratinocytes reveal horizontal dermokine gradient towards wound bed in non-healing wounds. (A)** Immunofluorescence images of patient tissue of healthy skin, well- and poorly- differentiated SCCs and psoriatic skin with the indicated antibodies. The scale bar for the anti-dermokine healthy skin image is 800 µm and for the anti-p120 image 500 µm. The scale bar for the well-differentiated SCC images is 1 mm, 500 µm for the poorly-differentiated SCC images and 250 µm for the images of psoriatic skin. **(B)** Magnified images from the squared area indicated in (**A**). The scale bar for the magnified images is 100 µm. **(C and F)**

Quantification of (**A**). Values are either raw Cy3 intensities (anti-dermokine (**C**)) or -p120 (**F**) of samples from at least N= 3 different patients. **(D)** Immunofluorescence staining of a non-healing wound bed (sample HS17.88) from a venous leg ulcer and the surrounding healthy skin. Scalebar = 1 mm (full image) and 100 µm (detailed view), respectively. **(E)** Immunofluorescence images of patient tissue of healthy skin and non-healing wounds with the indicated antibodies from the squared inset shown in Figure 5(A) The scale bar is 50 µm.
